# Non-canonical regulation of the plasma membrane copper transporter CTR1 through modulation of membrane mechanical properties

**DOI:** 10.1101/2025.03.28.645880

**Authors:** Subhendu K Chatterjee, Sumanta Kar, Siddhanta Nikte, Tisha Dash, Tanmoy Ghosh, Saptarshi Maji, Durba Sengupta, Bidisha Sinha, Arnab Gupta

## Abstract

We describe a non-canonical, membrane receptor-like regulation of the human copper transporter CTR1 in response to copper stimuli. CTR1 is the sole high-affinity trimeric plasma-membrane copper-importing channel that self-regulates by undergoing endocytosis to limit copper uptake. We observed that preceding copper-induced endocytosis, CTR1 forms clusters on the plasma membrane, a phenomenon that is typically observed in membrane receptors. We deciphered the mechanism of CTR1 clustering and studied its ramifications on plasma membrane physical properties harboring the clusters that favors endocytosis. Membrane tension and fluctuation are fundamental regulators of pre- and post-endocytic events. Using coarse-grain MD simulation and coupled Interference Reflection Microscopy-Total Internal Reflection Fluorescence Microscopy we demonstrated that CTR1 clusters induce positive membrane curvature, increase in local membrane tension and decrease in local membrane fluctuation; alterations that favors formation of endocytic pits. Clustering is facilitated by copper-sequestering Methionine rich extracellular amino-terminus of CTR1. MD-simulations and IRM-TIRF imaging revealed that CTR1 clustering is facilitated by membrane cholesterol, depletion of which delays CTR1 endocytosis. CTR1 clustering promotes clathrin-coated pit formation that engages recruitment of adapter protein AP-2. To summarize, we report a hitherto unknown ‘pre-endocytic’ ‘receptor-like’ phenomenon of ligand-induced clustering of a metal channel, that in-turn regulates self-endocytosis by modulating membrane properties.

## Introduction

Functions of ion channels and transporters are governed by a spectrum of mechanisms that include, ligand-gated, mechanically-gated, light-gated and cyclic-nucleotide gated regulations to name a few ^1–4^. Barring a few exceptions, conventionally channels function at the site of localization in the cell and are typically not translocated to other cellular locales in response to ligand binding or while transporting the ligand. In this study we report a very unique case where the high affinity copper transporter, CTR1 acts as a channel, importing copper and in the process gets endocytosed to limit excess copper uptake. We determined the mechanism of regulation of CTR1 by copper and how it affects plasma membrane (pm) properties making it primed to undergo endocytosis. Copper is a micronutrient essential for all eukaryotic organisms as it play crucial roles in coordinating different physiological activities of cells ^5^. First described in studies from Gitschier group using yeast complementation assays, the copper transporter CTR1 was implicated in copper import in mammalian cells ^6,7^. Cryo-EM studies revealed that CTR1 assembles as a trimer to attain a channel-like architecture ^8^. The authors demonstrated that the putative pore formed for copper ion translocation through the interface of the three identical monomers deviates significantly from the structural design of typical transporters and that copper uptake transporters have a novel ‘channel-like’ architecture ^8^. A second study from Unger’s group showed that the second transmembrane domain of the 3 TMDs of trimeric hCTR1 creates the ion-translocating pore that runs across the membrane bilayer at the interface between the three subunits ^9^. More recently, X-ray crystallographic structure of CTR1 at a resolution of 3Å from Atlantic salmon revealed that two layers of methionine comprised of M^150^ and M^154^ forms the selectivity filter, coordinating two bound Cu^+^ ions close to the extracellular entrance ^10^.

Copper shuttles between its two primary oxidation states, Cu(I) and Cu(II). Though participating in many physiological processes, the cuprous ion Cu(I) is unstable in the oxidizing environment; Cu(II) is the most abundant oxidation state in hydrophilic and oxidizing environments ^11^. We have previously shown that the human Copper Transporter-1 (hCTR1), besides importing copper, also plays a crucial role in maintaining the Cu(I) redox state that renders the metal bioavailable for physiological utilization in cells ^12^. The extracellular amino terminus containing multiple Histidine-Methionine (His-Met) motifs are key towards maintaining the Cu(I) redox state. Studies from others and our group have shown that hCTR1 localizes at the plasma membrane in copper chelated or basal copper (when copper is not added to the cell culture media) and shows strong colocalization with the pm marker Na,K-ATPase (*ATP1A1*). Interestingly, upon external copper treatment, CTR1 endocytoses to early endosomal compartments and common recycling endosomes (CRE) ^12,13^. Curnock and Cullen demonstrated that internalized CTR1 localises on retromer-positive endosomes and, in response to decreased copper, retromer regulates the recycling of CTR1 back to the pm ^14^.

Clathrin-mediated endocytosis (CME) is the major pathway for uptake of surface receptors and their bound ligands ^15–17^. CME is a complex phenomenon that is achieved through coordinated activity of multiple proteins that assist endocytic pit initiation, pit formation and ultimately scission of the clathrin-coated pits. The Adapter Protein, AP-2 is major regulator of CCPs. The AP2 complex links the PM–enriched phosphatidylinositol lipid, PI4,5P_2_, to the endocytic sorting motifs on cargo ^18–20^.

Besides synchronized participation of a spectrum of proteins, the membrane mechanical properties also play essential and concerted role to carry out a successful CME ^21^. Tension and fluctuations are the two major mechanical properties of a biological membrane that show inverse correlationship. Membrane tension in mammalian cells lie in the range of 0.001 – 3 mN/m ^22,23^. To operate within this stipulated range and maintain membrane homeostasis, mammalian cells adopt a large membrane area, that allows easy membrane fluctuations and formation of membrane reservoirs such as membrane folds, wrinkles, caveolae, vacuole-like dilations and blebs under low levels of tension ^24^. Since endocytosis requires invagination and internalization of patches of membrane from the pm, the tension at the sites of endocytosis is low. As endocytosis progresses, the positive membrane curvature generated by the CCP elevates the local tension and hence the local fluctuation drops.

Plasma membrane-residing ion channels do not typically endocytose in response to ligand binding or ligand transport. Rather receptor clustering is a common phenomenon and occurs when receptors on a cell surface are dispersed into localized clusters. It is a key feature of cell signaling and can affect how a cell responds to ligands. It has been previously demonstrated that receptors cluster at the plasma membrane upon ligand binding ^25^.In Clathrin Independent Endocytosis, using cytokine receptors, Salavessa et al calculated the critical number of receptors that should aggregate to induce endocytosis ^26^. A medium sized cluster composed of four to six receptors is key for IL-2R internalization and is promoted by interleukin 2 (IL-2) binding. Interestingly, larger clusters (> six receptors) are static and rather inefficient in internalization.

A great deal of evidence points towards the crucial role of inter-molecular interactions between receptors within a cluster play in transmembrane signalling. The robust systems exemplifying extensive and stable clusters are observed in the bacterial chemotaxis system ^27–30^ and the ryanodine receptors in excitable tissue ^31^. Generally, the adaptor proteins assist in the close localization and interaction between the cluster forming receptors. What is the functional ramification of such receptor clustering? Modulation of receptor-receptor interactions in a cluster is affected by the conformational states of adjacent receptors – i.e. whether they are active or inactive – can then cause conformational spread within a cluster. This conformational spread leads to ligand-binding to one receptor affecting the activity of a number of adjacent receptors in the cluster ^32,33^. As a consequence, clusters are instrumental in signal amplification by raising the sensitivity of receptors by more than an order of magnitude.

A few studies, primarily in neurons, have revealed that even ion channels exhibit clustering ^34–36^. Shuai et al ^37^ studied the distribution and clustering of inositol 1,4,5-triphosphate receptors in the plasma membrane that controls the flux of Ca^2+^ from the endoplasmic reticulum into the cytosol. By using mathematical modeling, authors show that channel clustering can enhance the cell’s Ca^2+^ signaling capability. Sato et al ^38^ provided a stochastic model of cluster formation of five different ion channel in the plasma membrane. They demonstrated that CaV1.2, CaV1.3, BK, and TRPV4 channels are randomly distributed on the plasma membranes of excitable cells and that form homogeneous clusters that grow in size until they reach a functional steady state. Ion channel clusters have been traditionally been thought to be causal to signal amplification and to propagate efficient downstream processes. Ohm et al showed that clusters of voltage-gated ion channels seen at the axon initial segment and the nodes of Ranvier trigger myelin formation ^36^.

In the present study we provide a view of a hitherto unknown mechanism of clustering of ion channel, not to engage signal amplification but rather as a self-regulatory mechanism to inhibit further ion uptake. We further delineate the mechanism of how these clusters affect mechanical properties of membrane that facilitates endocytosis of the channel clusters.

## Results

### Copper treatment promotes clustering of hCTR1 on the plasma membrane

CTR1 resides on the plasma membrane and imports copper, providing the required amount of copper to the cell. During elevated extracellular copper, CTR1 exhibits a unique ‘non-canonical’, membrane receptor-like regulation and undergoes endocytosis.

In order to comprehensively examine the distribution of CTR1 on the plasma membrane, we utilized Total Internal Reflection Fluorescence (TIRF) microscopy (Suppl. Fig.1A). We expressed mNeonGreen-tagged hCTR1 in HEK293 cells, fixed and imaged the cells under TIRF microscope. During basal copper condition, we noticed that small puncta of CTR1 were randomly distributed across the plasma membrane without any apparent localization pattern (Fig 1.A). As previously reported by us and other groups, we observed a decrease in the fluorescence signal intensity on the plasma membrane after exposing the cells to 100μM copper, indicating the Cu-induced endocytosis of hCTR1 (Fig 1. A) ^12,13,39^. Endocytosis, not being a common occurrence observed in channels and transporters, we were intrigued by this phenomenon of its regulation and delved further into it. The spatiotemporal dynamics of CTR1 endocytosis is rapid, and as a result, it is challenging to track the CTR1 units just before their disposal into Clathrin Coated Vesicles (CCV). In order to precisely investigate the pre-endocytic event of CTR1, we limited the generation of CCVs by exposing the cells to dynasore hydrate (Dh) (Fig 1. A); It inhibits dynamin family proteins in a reversible manner, which prevents the final scission of clathrin-coated vesicles. As a result, these immature vesicles stay as clathrin-coated pits on the plasma membrane. Incorporating copper treatment into Dh-treated cells allowed us to trap CTR1 units on the plasma membrane at the pre-endocytic stage. Remarkably, we observed the development of large CTR1 clusters across the plasma membrane with increasing duration of copper treatment. This suggests that, upon Cu-induced endocytic signal, randomly distributed CTR1 units are directed to form clusters, resulting in formation of endocytic patches (Fig 1. A). We quantified and compared the hCTR1 signal intensities at pixel level under various circumstances. During the pre-endocytic stage, we observed the presence of considerably large cluster in copper treatment as compared to the basal condition. We compared and analyzed the size distribution pattern of hCTR1 prior to clathrin-mediated endocytosis (CME) in copper treated and control conditions (Fig 1. A; right panel).

**Figure: 1.**
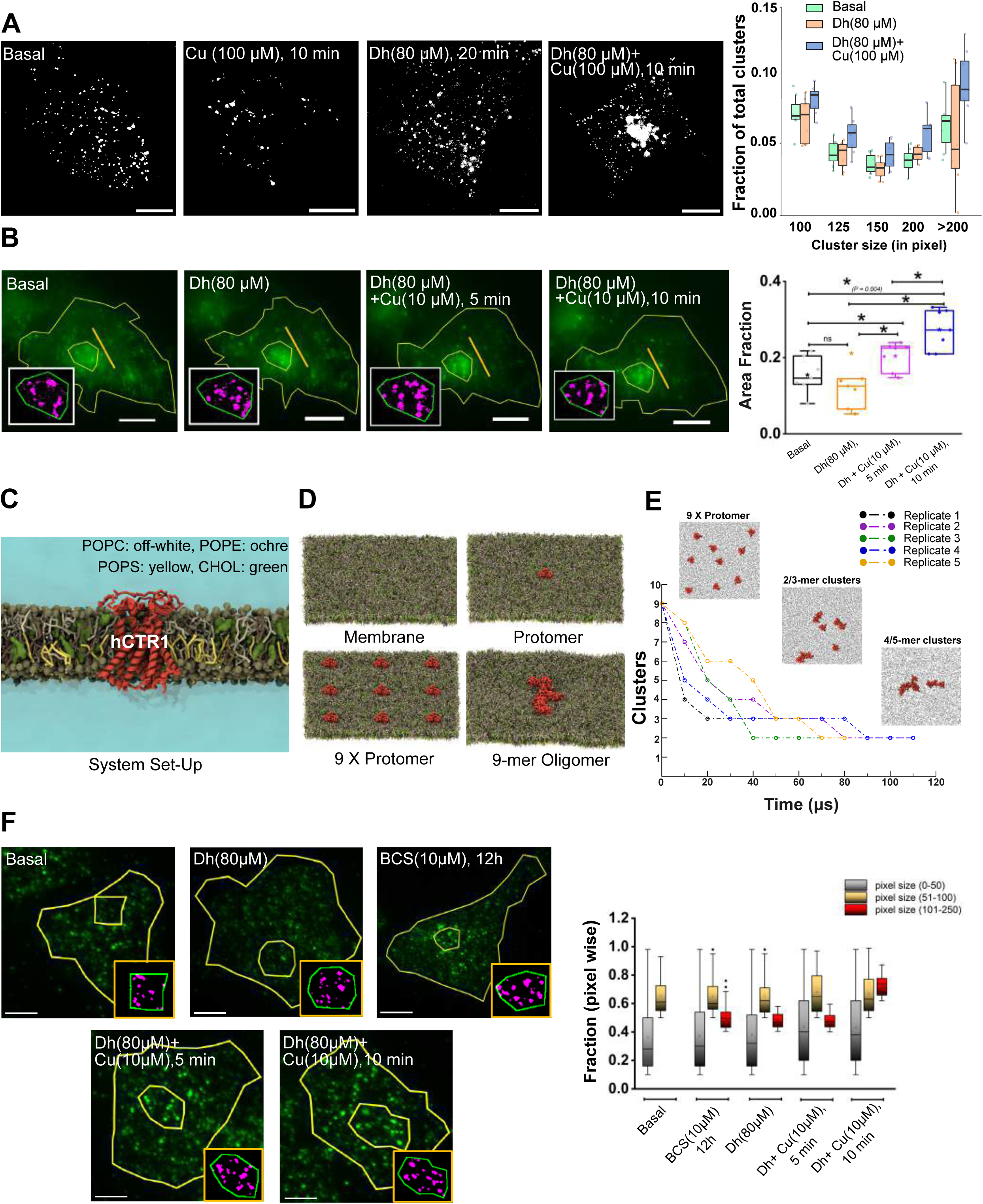
Clustering of the copper transporter CTR1 affects its self-regulatory endocytosis. A. Fixed cell TIRF imaging of FLAG-hCTR1 at basal, copper (100 µM), dynasore (Dh; 80 µM, 20 min) and Dh+Cu (100µM for 10 min) conditions (from left to right). The graph (right) exhibits the fraction of cluster size of FLAG-hCTR1 at basal, dynasore (Dh; 80 µM, 20 min) and Dh+Cu (100µM for 10 min) conditions (images were derived from TIRF and analyzed in ImageJ^Ⓡ^). B. Representative TIRF image of FLAG-hCTR1 in transfected in the same HEK293T cell for basal, dynasore (Dh; 80 µM, 20 min) and Dh+ Cu (10µM; for 5 &10 min) respectively (*from left to right*). In the zoomed-inset, the same ROI was processed using MATLAB R2020a in the different treatment conditions mentioned above. The graph (right) is plotted for comparison of area fraction (Area covered by puncta/Total area of ROI) among the treatments mentioned above. Statistical analysis was performed using Mann–Whitney U-test, ns denotes P value>0.05. *denotes P-value<0.05. Scale bars: 10 µm. C. System snapshot of trimer unit of hCTR1 i.e. protomer (tan red) in model membrane. The lipids are shown in colors as; POPC: off white, POPE: ochre, POPS: yellow, Cholesterol: green, lipid head groups in vdw representation in greyish green. D. System snapshots represent the model membrane (left upper panel), protomer in the membrane (left lower panel), 9x protomer in the membrane (right upper panel) and 9-mer oligomer in the membrane (right lower panel). E. Cluster formation of hCTR1 over simulation time, at time points in the range of 10 microseconds for 5 replicates. The snapshots of the observed hCTR1 protomer clusters and their sizes are shown in the above panel. Protomer shown in red color and only lipid headgroups are shown in black color for clarity. F. Representative TIRF images of endogenous hCTR1 labeled with anti-CTR1 antibody in HEK293T cells for basal (untreated), copper chelator, bathocuproinedisulfonic acid disodium salt (BCS; 10µM for 12 h) treated, only Dh treated, Copper (10µM; for 5 and10 min preincubated with 80µM Dh). Graph (right) compares the fraction of puncta sizes grouped as 0-50, 51-100 and 101-250 pixels under the abovementioned treatment conditions..

We spatiotemporally tracked the growth pattern of CTR1 clusters from the basal condition to Cu-treated condition. This allowed us to track a specific CTR1-expressing region of interest (ROI) on the cell surface under various stages of excess extracellular copper conditions. Observations revealed that initially dispersed hCTR1s clustered approximately 5 minutes after initiating a 10μM copper treatment in the presence of Dh. Cluster formation became more pronounced after 10 minutes, (Fig 1. B, left to right; inset LUT images highlighting the “objects” (clustered pixels) detected at regions of hCTR1 expression). On quantifying the area fraction (fractional area covered by the puncta detected as objects), the increase of footprint of hCTR1 puncta was recorded, due to the cluster formation of hCTR1 on pm prior to clathrin-mediated endocytosis (Fig 1. B, right panel bar plot).

Protein clusters are biophysical entities on the pm with defined mass and size. We hypothesized that CTR1 clusters modulates membrane properties that engages endocytic pit formation. Towards that goal, we performed coarse grained simulations of the functional trimeric unit of hCTR1, i.e., protomer(s) embedded in the lipid bilayer (Fig 1C). We varied the number of protomers in the lipid bilayer with systems such as protomer (single channel) and oligomer (multimers) states. Representative snapshots of the systems are shown in (Fig 1D). All the hCTR1 protomer units remain embedded in the lipid bilayer during simulations and diffuse freely. Several protein-protein contacts were sampled, and protein multimers were observed. Here, we report the membrane perturbations observed in the oligomeric (1 cluster of 9-mer oligomer) state as well as other small clusters observed during the multimer simulations compared to only membrane and protomer (1x protomer) systems, highlighting the effect of cluster on the membrane.

Initially, to understand hCTR1 inter-protein interaction and clustering, we analyzed the hCTR1 protein cluster formation in a multimer (9x protomer) system over simulation time for each replicate. We observe the interaction and diffusion of the hCTR1 proteins in the membrane and their subsequent formation of larger clusters of different sizes (i.e., hCTR1 protomer units in one cluster) at different timescales during simulation time, shown in Fig 1E. We observe the initial hCTR1 protein interactions and small-size (i.e., 2/3-mer cluster) cluster formation at 6μs simulation time, eventually forming 2 clusters with a size of 4/5 hCTR1 protomer in each, i.e., 4/5-mer cluster. In all the replicates, we observe two clusters of hCTR1 proteins at the end of the simulations, even at a timescale of 100μs. Once these clusters are formed, they remain intact, and we do not observe any dissociation of hCTR1 protein from clusters during the simulation. The protein interaction and formation of stable clusters of hCTR1 observed in simulations indicates, and as confirmed by the experimental results of TIRF, that hCTR1 does have an inherent propensity to interact and form clusters and puncta. Further, we performed biased simulations to form larger 9-mer oligomers that remained stable in the timescales of the simulations (Figs 1D and 1E). To mimic the simulations and to test if CTR1 has an inherent property to form clusters, we compared endogenous CTR1 cluster formation in copper-depleted (prolonged copper-chelator, BCS treatment) versus copper treatment in presence of Dh, studying role of copper in CTR1 cluster (Fig 1F, left panel, images). Quantitative analysis of area fractions and puncta pixel size indicated that CTR1 clustering may be a constitutive phenomenon, but it is not sufficient to trigger its CME (Fig 1F, right panel). High copper promotes and enhances clustering, triggering endocytosis of CTR1. (Figs 1A)

### CTR1 clusters induce physical alterations in Plasma Membrane properties that favours Endocytosis

Based on findings from the previous section we hypothesize that copper-mediated CTR1 clustering favours formation of endocytic pockets. Previous studies have established that endocytosis is influenced by membrane tension, curvature and membrane elasticity ^40–42^. We tested whether clustering of hCTR1 alters membrane properties that favours initiation of endocytosis.

We used the afore-mentioned set of simulations to investigate membrane perturbations observed around the hCTR1 protein clusters. To understand the effect of protein on the overall lipid bilayer, we first calculated the lipid order parameter. The lipid order parameter reports on the flexibility of the acyl chain of the lipids ^43^. We determined the lipid order parameter for all the lipid types in the simulations system in five independent replicate runs (Suppl. Fig. 2A). We observe a decrease in the lipid order in both chains of lipid types in the hCTR1 oligomer state compared to the protomer and the only membrane system. We also observed a decrease in lipid order around 2nm of hCTR1 protein compared to the bulk lipid order parameter. Further, when systems were compared, it was observed that membrane thickness was reduced around the oligomer more than the membrane control and protomer system (Suppl Fig. 2B). For the last 5μs of multimer systems, we noted a significant reduced thickness around clusters of hCTR1 proteins. This suggests that the hCTR1 clusters affect the lipid flexibility and lipid bilayer thickness at the local membrane region suitably priming the cluster sites for endocytosis.

Based on the membrane alterations observed around hCTR1 clusters we hypothesize that these plasma membrane changes engage in formation of the endocytic pit that internalizes CTR1 in response to copper. Endocytic pit formation is coupled with changes in membrane curvature ^44^. We calculated the membrane curvature for all the systems. We did not observe any appreciable curvature in the model membrane, protomer, and in the membrane devoid of the hCTR1 protomer clusters, suggesting that they maintain flat surfaces. However, for the 9-mer oligomer and small clusters (2/3-mer and 4/5-mer clusters) in multimer systems, we observe the formation of the positive curvature in a membrane leaflet around these clusters (Fig. 2A; replicates are shown in Suppl Fig. 3A). However, the curvature observed around 9-mer oligomer is more prominent than the curvature observed around the small clusters. These findings suggest that the hCTR1 cluster formation induce curvature in the membrane as illustrated in Suppl Fig. 3B.

**Figure: 2.**
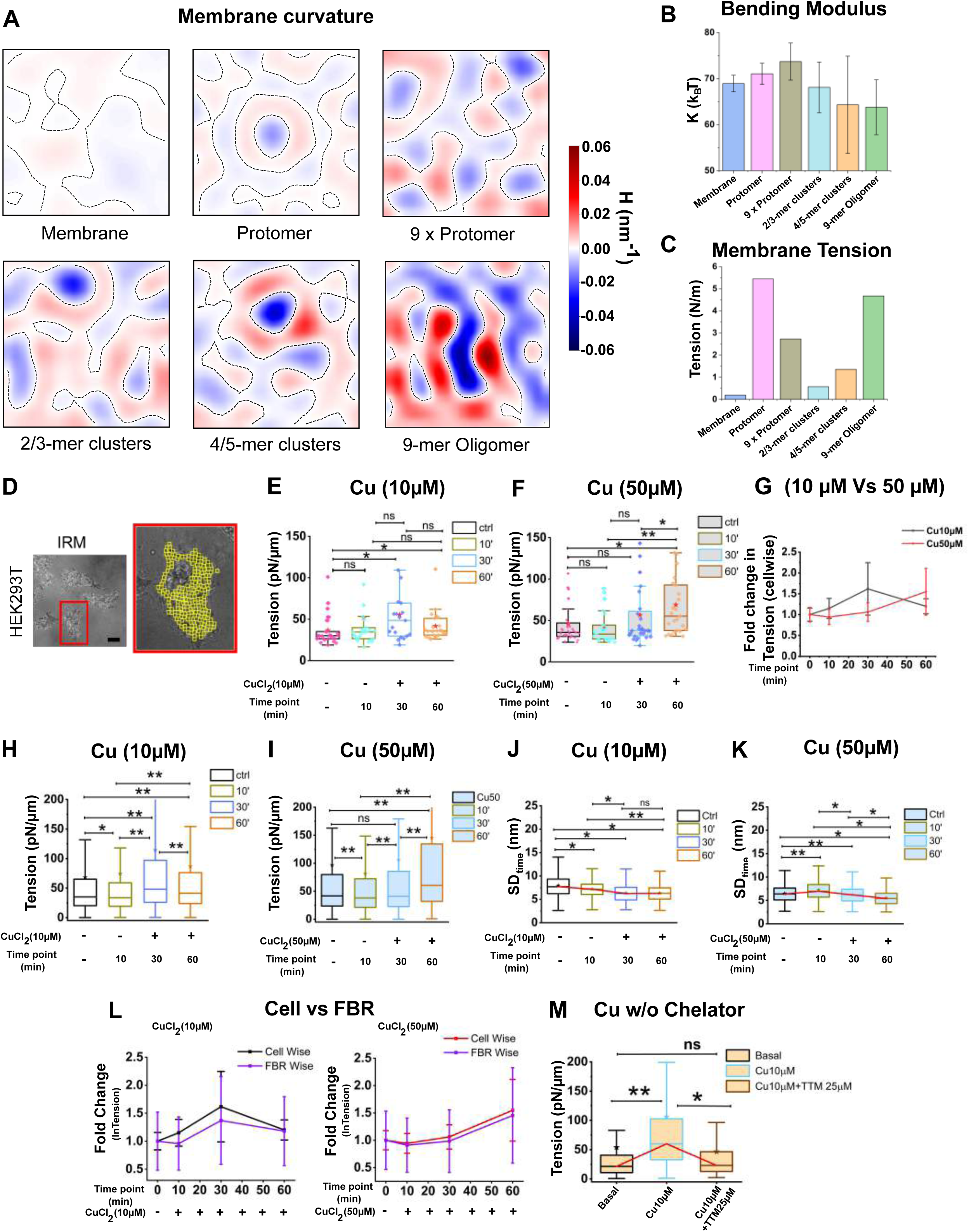
Clustering of hCTR1 alters plasma membrane mechanical properties. A. 2D map represents the membrane curvature observed in the lower leaflet of the membrane during different phases of hCTR1 cluster formation in the MD simulations. Representative plots are shown for the systems with only membrane, protomer in the membrane, 9x protomer in the membrane, 2/3-mer clusters, 4/5-mer clusters, and 9-mer oligomer in the membrane. B. The membrane bending modulus for the above-mentioned systems is plotted as a bar plot with standard error calculation across all replicates. C. The membrane tension is plotted as a bar plot for the above-mentioned systems simulated as independent simulations. D. Representative DIC images (left panel) and IRM images (right panel) of HEK293T cell; maximize view of the cell shows the FBR regions as small yellow boxes, Scale bars: 10 µm. E& F. Cell-wise tension profile for Basal and Copper (10µM & 50µM, 10 min, 30 min and 60 min, respectively, from three independent experiments. Where *‘n’* cells (for 10µM) = 25, 26, 19 & 23 and *‘n’* cells (for 50µM) = 31,23, 28 & 25 respective to basal and different copper conditions. from three independent experiments. G. Comparison between cell-wise fold change in tension for 10 µM and 50 µM copper treatment at earlier mentioned different time-lags. H & I. FBR-wise tension profile for Basal and Copper (10µM & 50µM, 10 min, 30 min and 60 min, respectively, from three independent experiments. Where *N*_FBRs_ (for 10µM) = 3911, 3499, 3712 & 3030 and *N*_FBRs_ (for 50µM) = 4394, 3840, 2927 & 3821 respective to basal and different copper conditions. from three independent experiments J & K. Comparison of Spatial fluctuation [SD _time_] between Basal and Copper (10µM & 50µM, 10 min, 30 min and 60 min, respectively, from three independent experiments. L. comparison between cell-wise Vs FBR-wise fold change in tension for basal and copper (10 µM and 50 µM) treatment at earlier mentioned different time-lags. M. FBR-wise tension profile in mNeon hCTR1 transfected HEK293T cell. Where *N*_FBRs_ = 385, 310, 318 respective to basal, Cu10µM, 10 min followed by treatment with 10 min potent copper chelator, Tetrathiomolybdate, (TTM; 25 µM). Statistical analysis was performed using Mann–Whitney U-test, ns denotes P value>0.05. *denotes P-value<0.05** denotes P-value<0.001. Scale bars: 10 µm.

Further, to probe further into membrane bending, we calculated the bending modulus of the membrane, revealing how oligomerization impacts membrane elasticity. The bending modulus of a membrane is a physical constant that measures the amount of energy required to change the curvature of a membrane from its normal state The bending modulus, is a direct correlation of the membrane’s resistance to bending, varies significantly under different conditions (Fig. 2B). In the absence of hCTR1 protein in the membrane or in the presence of a single protomer (1x protomer), the bending modulus of the membrane ranges between 68 kBT to 72 kBT, indicative of the inherent stiffness of the membrane. When more protomers are present, the bending modulus increases to about 74 kBT, suggesting increased stiffness due to the large number of protomers. However, concurrent with increase in membrane fluidity, once the protomers start to cluster, we observe dimers/trimers or tetramers in the membrane, the bending modulus decreases to 68 kBT. When the final larger 9-mer oligomeric cluster is formed, the bending modulus reduces to 63 kBT. The decrease in the bending modulus with increasing clusters is concurrent with the increased membrane curvature observed with the onset of oligomerization. Our analysis shows that prominent curvature formation occurs in oligomer state simulations, corresponding to the reduced bending modulus as compared to multimer simulations with no clusters to a few small hCTR1 clusters, protomer, and membrane alone. These findings highlight the complex interplay between protein oligomerization and membrane mechanical properties, providing insights into the functional dynamics of membrane proteins and membrane-associated processes.

We calculated membrane tension for the different systems of hCTR1 proteins in the membrane (Fig. 2C). We observe the lowest membrane tension in the absence of protein and the highest in the presence of a protomer state. Surprisingly, when multiple protomers are present, the membrane tension is reduced. Further, membrane tension reduces when the protomers start to cluster, i.e. with small dimers/trimers or tetramers clusters of hCTR1. However, in the presence of the oligomer, we observed an steep rise in membrane tension.

### CTR1 clusters modulates local membrane tension and fluctuations *in vivo*

Encouraged by our coarse-grained analysis, we sought to understand the role of CTR1 clusters in modifying plasma membrane properties using *in-vivo* experimental approaches. We utilized IRM (Interference Reflection Microscopy), a non-invasive imaging technique, to assess the physical characteristics of the basal membrane during various stages of hCTR1 endocytosis mediated by copper. In IRM, interference patterns are caused by the dual reflection of light – at the coverslip-media interface and the basal plasma membrane both occurring due to the differences in refractive indexes of the cell basal membrane and the glass coverslip from that of the media (Suppl. Fig. 4A). The intensity obtained in the IRM images of the basal cell membrane (Fig.2D) were converted to relative height using calibration steps performed using a 60 μm polystyrene bead (Figs Suppl Fig 4B). Calibration was applied on specific regions termed as First Branch Region (FBR) (typically membrane regions that are within ∼ 100 nm of the coverslip) for all analysis. The time-series of the relative height of the basal membrane reflects the height variations or membrane fluctuations. The amplitude of the variations was captured by the parameter (SD_time_ – standard deviation of height fluctuations in time). The time-series were also utilized to also subsequently derive cell surface tension using the methodology described by Biswas and co-workers ^45^ and also elaborated in the methods section.

Measuring fluctuations and tension in cells (HEK293 expressing endogenous CTR1, marked by red box in Fig. 2D), we observed that with low copper treatment (10 μM), relative to the basal state, average cellular tension in cells increased mildly at 10 min. By 30 min, tension peaked, suggesting the occurrence of a ‘membrane tug’ that leads to endocytosis. After this 30-minute point, tension started to decrease, indicating the completion of endocytosis and the return of the membrane to a homeostatic state by recycling (Fig 2E). Contrastingly, in high copper treatment (50 μM), we noticed an intriguing pattern. Till the 10-minute time point, membrane tension initially dipped but then gradually increased over time (Fig 2F). We argue that at a higher copper treatment, the pm primes itself for sustained endocytosis and hence adds membrane to bring about successful prolonged endocytosis, leading to increased fluctuations and lowering of tension. The initial reduction in tension at a 50 μM treatment may reflect the interplay of two competing processes influencing to tension as suggested by predictions in Fig 2C. Considering the simulation-based prediction of tension reduction during early stages of clustering (Fig. 2C – comparing protomer to cluster), we believe the initial decrease in likely due to the abundant formation of pre-clusters at 50 μM that lower tension, which are subsequently converted into oligomers and tension-enhancing pit-shaped structures. The comparative analysis fold change in membrane tension in high and low copper along the time scales supports our hypothesis (Fig. 2G).

We next focused on analysing the tension change in the plasma membrane by measuring tension and fluctuation-amplitude at local FBRs (Fig.2D, right panel marked by yellow boxes). The FBR-wise pooled tension measurements, for both 10 μM and 50 μM copper treatments showed a similar tension profile at early time points, suggesting that the effects of copper on membrane properties are also locally manifested (Fig 2H and 2I). Sustained high and increasing membrane tension in cells treated with higher copper (50 µM) indicates continued pinching-off of endocytic vesicles possibly harbouring CTR1. The amplitude of temporal height fluctuations, (SD_time_) – unlike tension, is a model-independent measurement, and also displayed the same (Fig. 2J and 2K) – effect of concentration and treatment time on membrane dynamics – with higher fluctuations in cells with lower tension and *vice versa*. A comparative analysis of tension fold change of whole cell and FBR reveals a similar trend in both high and low copper treatments (Fig 2L). This comparative analysis also essentially indicates that in conditions of ‘induced endocytosis’, i.e., copper-induced endocytosis, the phenomenon majorly overrides the membrane properties over ‘constitutive endocytosis’.

When cells were treated with the copper chelator TTM (25 μM) following copper exposure, FBR-wise tension profiles returned to normal, indicating that copper-induced changes in membrane properties are likely linked to CTR1 endocytosis in high copper conditions, a hypothesis we explore further in our investigation (Fig 2M).

This phenotype in low copper treatment demarcates a gradual increase of membrane tension due to Cu-mediated endocytosis of hCTR1 up to a certain time point (30 min) and signifies vesicular recycling back to the PM that contributes to membrane addition at the PM which is well reflected as the membrane tension falls at later time point. However, during high Cu (50 μM) treatment, we observed a trend where the initial membrane tension dipped due to cells maintaining the ion homeostasis through a rapid rate of hCTR1 endocytosis and recycling at that time point. It is noteworthy that the synchrony between CTR1 endocytosis and membrane tension plays a significant role in maintaining the surface distribution of CTR1 and hence, copper uptake.

### The plasma membrane abundance of CTR1 impact local membrane properties

We aimed to understand the sequence of events of CTR1 endocytosis upon Cu treatment and how membrane modulations contribute to it. Using correlative live IRM-TIRF imaging, we co-assessed the membrane alterations and CTR1 abundance on the plasma membrane. mNeonGreen-tagged CTR1 was expressed in HEK293 cells, and the readouts were analyzed spatially and temporally emphasizing on local region (FBR), harboring the clusters.

It is important to mention that we identified both low and high exogenous mNeon-CTR1 expressing zones in a single cell (Fig 3. A & B). Magnified TIRF images of high expression (larger inset) and low expression zones (small inset) in a single cell and their respective heat-maps representing the abundance of CTR1 pre- and post-copper treatments is shown in the lower panels of Fig.3 A & B. This provides us with a convenient tool for investigating whether CTR1 expression levels at the pm influence its endocytic self-regulation. Without any external copper treatment (Basal condition), significant difference in fluorescent intensity distinguish high CTR1 expressing zones (HE zone) from low expressing zones (LE zone) (Fig 3. C, left bar plot). Surprisingly, we observed that these two zones have comparable levels of membrane tension in absence of external copper treatment (Fig3. D, left bar plot). This suggests that regions with high levels of CTR1 expression do not have particularly high membrane tension, which means that they do not provide any favourable conditions for endocytosis in absence of external copper treatment, that aligns well with our finding described in Fig.1F. Following a 10-minute treatment with 10μM Cu, we observed a slight reduction in intensity indicating initiation of endocytosis (Fig 3. C, right bar plot). Remarkably, when compared to the LE zone, the HE zones exhibited notably elevated membrane tension, indicating a greater degree of endocytosis (Fig 3. D, right side bar plot). Further comparing the fold change in tension before and after copper treatment shows a higher increase in the HE zones compared to the LE zone (Fig 3. E). This result suggests that during Cu-induced endocytosis, HE zones of CTR1 experience faster and more intense endocytosis. These findings substantiate the existence of variations in the expression levels of CTR1 coupled with copper levels modulating membrane properties (shown in Fig 3. F).

**Figure: 3.**
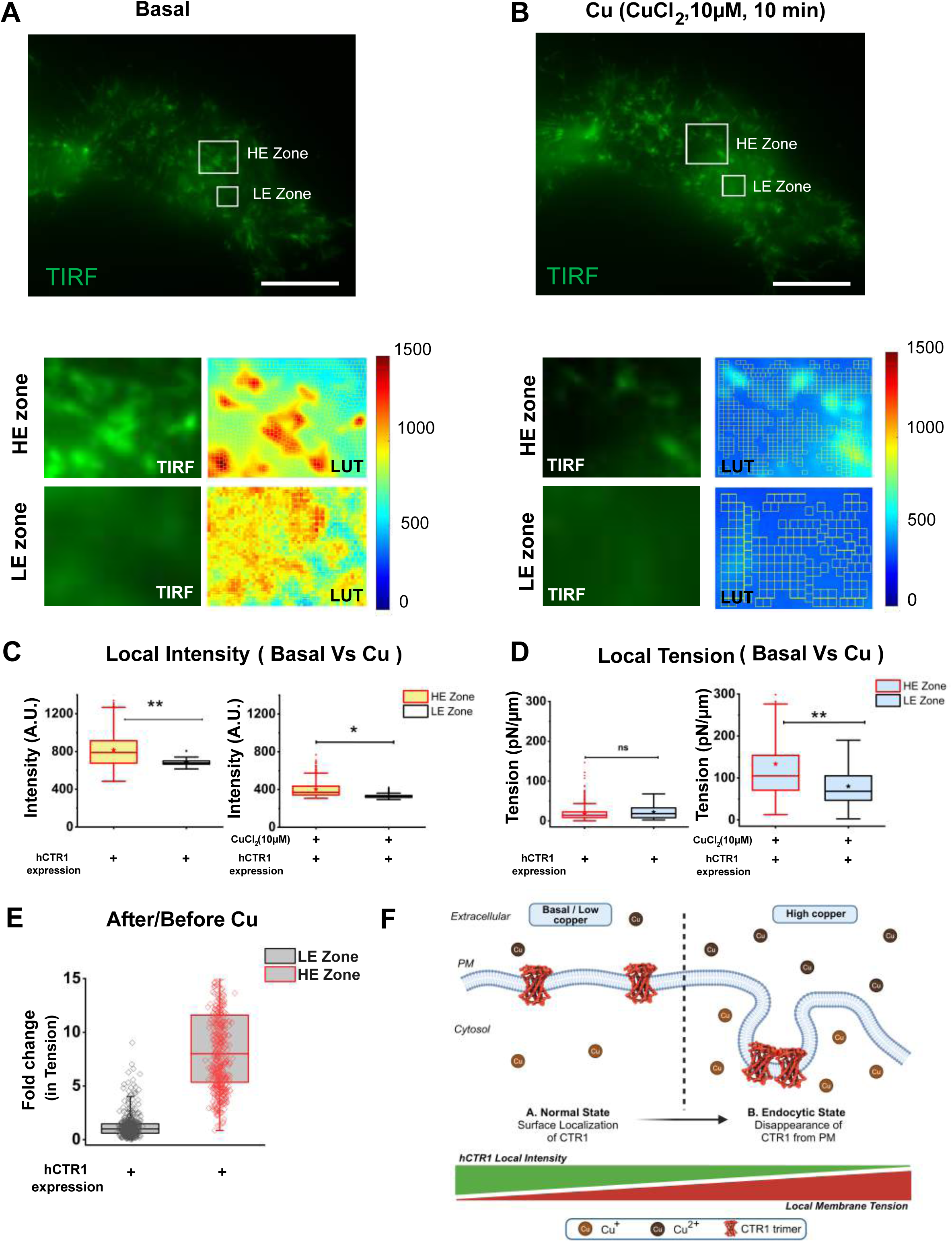
Local membrane tension correlates with local plasma membrane abundance of CTR1. A & B. Representative TIRF image of a single HEK293T cell at basal condition **(A)** and then treated with copper **(B)**. Bottom panels shown magnified view for high CTR1 expressing High expression (*HE*) and Low expression (LE) zones labeled as rectangular box and the corresponding heat map (*Lookup table*) against intensity for the same zone side by side. C. Comparison of local hCTR1 fluorescence intensity between HE and LE zone for basal (left side bar plot) and excess extracellular copper (10 µM; for 10 min) exposure (right side bar plot). *No. of FBRs* = 36 & 22 for HE and LE zones respectively). D. Divergence of FBR-wise local tension profile between HE & LE zone for Basal (left bar plot) and above-mentioned copper condition (right bar plot). *No. of FBRs* = 36 & 22 for HE and LE zones respectively). E. Differences in fold change ration (between after and before copper treatment) of normalized tension profile at HE and LE zone. Statistical analysis was performed using Mann–Whitney U-test, ns denotes P value>0.05. * denotes P-value<0.05** denotes P-value<0.001. Scale bars: 10 µm. F. Model depicting relationship between local state of hCTR1 and membrane property modulation at excess extracellular copper conditions (Created in BioRender. Gupta, A. (2024) BioRender.com/z49w299).

### Copper treatment enhance CTR1 clustering and subsequent endocytosis

So far, we have pinpointed plasma membrane tension and CTR1 clustering as two interconnected key regulators of CTR1 endocytosis. We asked whether the link between CTR1 clustering and modulations in membrane tension is regulated by external Cu.

In order to observe the pre-endocytic CTR1 clustering and test any accompanying changes in membrane characteristics, we employed correlative TIRF-IRM and monitored a specific region of interest (ROI) under various copper-induced endocytic conditions in presence of dynasore (Basal and Dh± Cu) (Fig 4. A, upper panel, left to right). We found that Cu triggered the clustering of CTR1 at the 5 min and 10 mins time points in the pre-endocytic state which reflected in a significant increase of intensity against hCTR1 (Fig 4A, upper panel). We compared two inversely inter-dependent membrane properties-fluctuation and membrane tension in basal and 10 µM Cu treated conditions. We observed a subsequent decrease in membrane fluctuation following an initial increase (Fig 4. A second middle panel). Exact opposite trend was detected in case of pixel-wise mapping of membrane tension in similar conditions (Fig 4. A, third middle panel) respectively. The R2 map shows significant association between tension and membrane properties use for the pixel-wise mapping of membrane tension (Fig 4. A, lower panel).

**Figure: 4.**
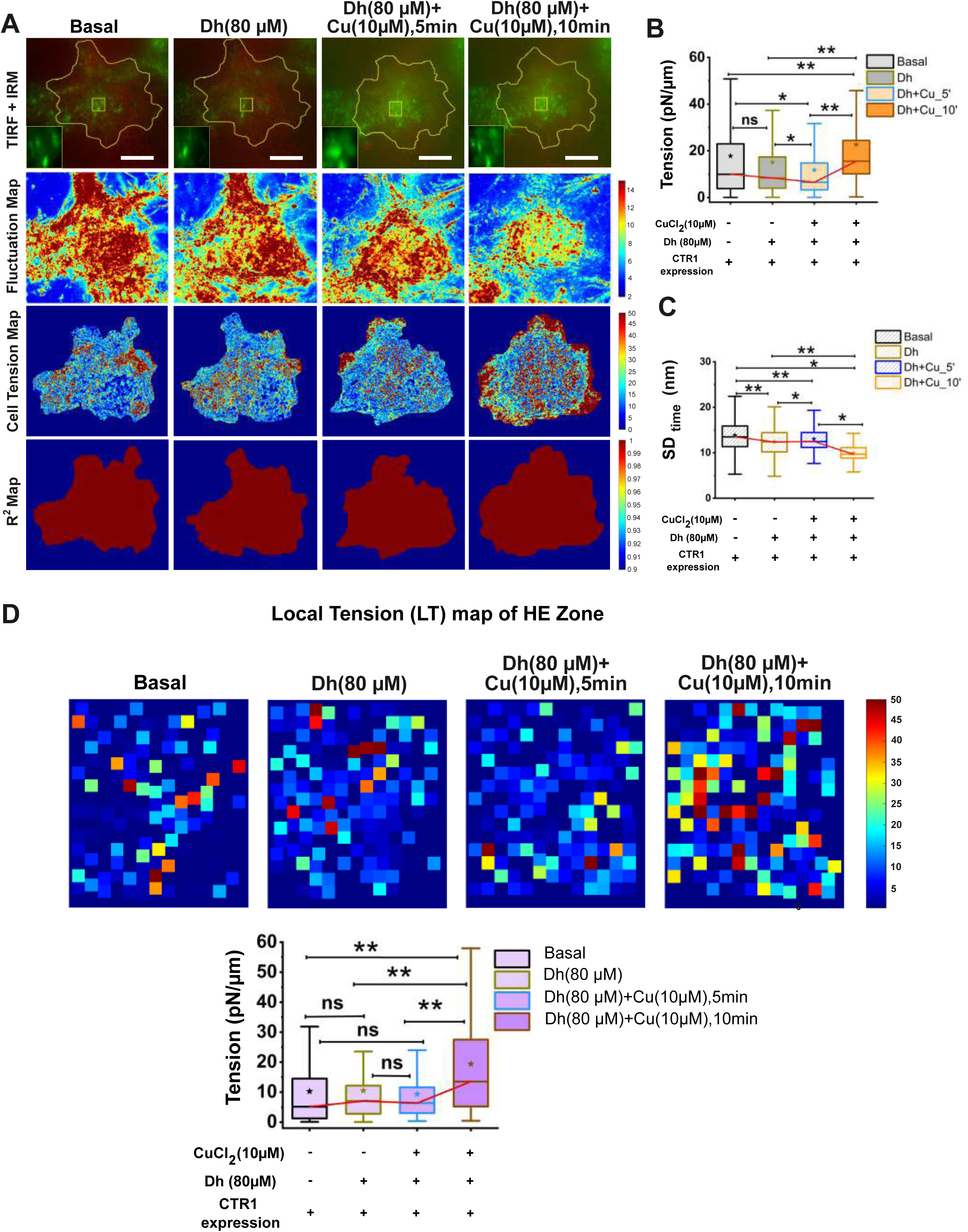
Correlation between copper-induced hCTR1 clustering and modulation of membrane mechanics. A. Upper panel (*Left to right*): Representative TIRF image of mNeonGreen-CTR1 of same HEK293T cell merged at basal, dynasore (Dh; 80 µM), Dh+ Cu (10µM; 05 min and 10 min). Zoomed-in view at left lower corner showing local high hCTR1 expression (*HE*) zone]. 2^nd^ panel and 3^rd^ panels (*Left to right*) show fluctuation map & tension map (pixel wise) respectively of the same cell at above-mentioned treatment conditions. Lower panel represents the R^2^ map of the same cell at above-mentioned treatment conditions. B. FBR-wise tension profile comparison between basal, dynasore (Dh; 80 µM), Dh+ Cu (10µM; 05 min and 10 min) treatments of the same cell. *No of FBRs* = 253, 248, 169 & 156 for basal, dynasore (Dh; 80 µM), Dh+ Cu (10µM; 05 min and 10 min) respectively. C. Comparison of spatial fluctuation (SD _time_) of same FBRs against different copper treatments mentioned above (where *No. of FBRs* are = 253, 248, 169 & 156 respectively). D. *Upper panel* represents a heat map against local intensity for the high hCTR1 express zone (*HE,* for the same zoom-in region marked as the yellow rectangle box in the upper panel under Fig.4A. among above mentioned different copper treatments (*from Left to right*). *Lower panel* shows the bar graph comparing local tensions of HE zone against same treatment conditions. Statistical analysis was performed using Mann–Whitney U-test, ns denotes P value>0.05. * denotes P-value<0.05** denotes P-value<0.001. Fluctuation map, pixel wise tension map and Local tensions vs local fluorescence intensity was plotted using MATLAB (R2020a). Scale bars: 10 µm.

Quantitative analysis showed slight reduction in cell-wise tension at the initial ‘pre-endocytic’ state (Dh + Cu,5min) signifying the initiation of clathrin-coated pits (CCPs) formation whereas a significant increase in tension at the later time point (Dh + Cu,10min) indicates tug in effective plasma membrane possibly due to its utilization in elongation and maturation of CCPs (Fig 4. B). At the same time point, spatial fluctuations (SD_space_) showed the opposite trend (Fig 4. C) as we illustrated earlier in Fig 4. A.

In continuation with the pre-endocytic state, we focused on comparing hCTR1 localization and membrane tension alteration following the same local region of interest (ROI) marked in a box in Fig. 4A top panel. The ROI was selected at a high expressing CTR1 zone (HE). We found a similar membrane tension scenario in ROI as we found in a cell-wise manner. Interestingly if we counter these observations with the tension map (Pixel wise) for the same time point, we found a slight reduction of tension at the initial time point (5 min) followed by a significant increase in tension at 10 min copper treatment (Fig 4D, left pixel-wise heat map and Fig.4D right bar plot).

### The pre-endocytic CTR1 clustering is dependent on its copper-sequestering amino terminal domain

We and others have previously reported that the amino-terminal of CTR1 harbors conserved Methionine motifs that are crucial in CTR1 endocytosis and copper import ^12,46^. We aimed to identify whether the Met rich amino-terminus facilitates copper-induced clustering of CTR1. Friedemann et al ^47^ demonstrated that the Methionine residue on the Aβ peptide, which is responsible for generating Alzheimer’s disease, plays a significant role in the aggregation of the peptide, and this involvement depended on the presence of copper. Taking into account the findings from this study, we tested the involvement of crucial amino-terminal methionine residues (M7GM^9^, the first Met-motif and ^40^MMMMPM^45^, the second Met-motif) of CTR1 in the genesis of clusters. These two distinct methionine patches are instrumental for the copper import activity of the transporter ^12^. We studied two distinct hCTR1 mutants; the first mutant, ΔM1M2-hCTR1, lacks both methionine patches, while the second mutant, Δ30-hCTR1, first 30 amino acids truncated, thus devoid of the first Methionine patch. We assessed Cu-induced cluster formation in these mutants and compared it to wild-type hCTR1 (mNeonGreen-hCTR1). In wild type hCTR1, the minimum average distance between puncta decreased significantly with extended copper treatment indicating the fusing or close apposition of puncta during same ROI follow up (Fig 5A & B). However, the distance between puncta in two CTR1 mutants showed no significant changes during copper treatment (Fig 5C,5D and Fig. 5E, 5F). Fig 5. G illustrates a comparative examination of cluster formation, represented as the area fraction on the plasma membrane of the hCTR1mutants and the wild type. For ΔM1M2-hCTR1, no such significant increase in cluster formation (area fraction) during the first 30 minutes of Cu treatment was recorded. Similarly, the Δ30 mutant did not exhibit a substantial increase in cluster formation under Cu-treated conditions. These results indicate that the intact amino terminus of hCTR1 containing the Met residues is crucial for clustering, which we establish as an essential pre-endocytic step that regulates subsequent endocytosis of CTR1.

**Figure: 5.**
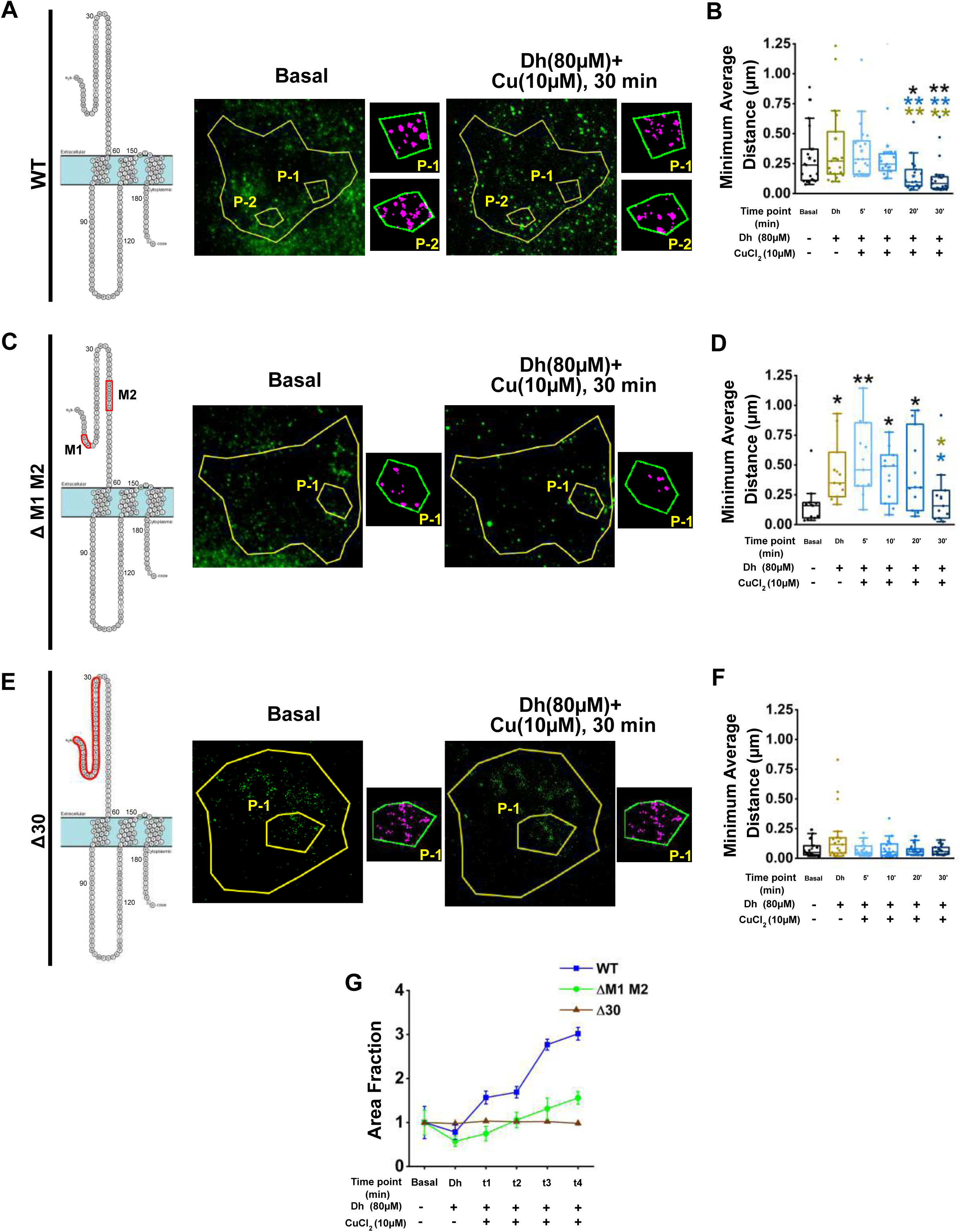
The copper-binding amino terminus of CTR1 facilitates its clustering on the plasma membrane. (A) Cartoon depicting different domains of wt-hCTR1 protein. Cells expressing wild-type mNeonGreen-CTR1 were treated with copper in presence of Dh and TIRF imaging of live cells was conducted at 05 min, 10min, 20min and 30 min. Representative TIRF image (pseudo color, green) of the same cell at basal and post-30min copper treatment. Zoomed regions showing two ROI -P1 & P2. (B) Comparison of minimum average distance (μ) between two nearby hCTR1 puncta of the ROIs against basal, dynasore (Dh; 80 µM, 20 min), Dh+ Cu (10µM; 05 min,10min, 20min and 30 min) conditions tracked in the same cell. *n_ROI_*= 21 (C) Cartoon depicting the mutant ΔM1M2-hCTR1 protein. Cells expressing mNeonGreen-ΔM1M2-hCTR1 were treated with copper in presence of Dh and TIRF imaging of live cells was conducted at 05 min, 10min, 20min and 30 min. Representative TIRF image (pseudo color, green) of the same cell at basal and post-30min copper treatment. Zoomed region shows ROI -P1. (D) Comparison of minimum average distance (μ) between two nearby hCTR1 puncta of the ROIs against basal, dynasore (Dh; 80 µM, 20 min), Dh+ Cu (10µM; 05 min,10min, 20min and 30 min) conditions tracked in the same cell. *n_ROI_*= 11 (E) Cartoon depicting the mutant Δ30-hCTR1 protein. Cells expressing mNeonGreen-Δ30-hCTR1 were treated with copper in presence of Dh and TIRF imaging of live cells was conducted at 05 min, 10min, 20min and 30 min. Representative TIRF image (pseudo color, green) of the same cell at basal and post-30min copper treatment. Zoomed region shows ROI -P1. (F) Comparison of minimum average distance (μ) between two nearby hCTR1 puncta of the ROIs against basal, dynasore (Dh; 80 µM, 20 min), Dh+ Cu (10µM; 05 min,10min, 20min and 30 min) conditions tracked in the same cell. *n_ROI_*= 19 Representative images (A, C and E) were modified from Protter. Statistical analyses (B, D and F) were performed using Mann–Whitney U-test, ns denotes P value>0.05, * denotes P-value<0.05, ** denotes P-value<0.001 (where black color asterisk for Basal Vs treated, grey color asterisk for Dh Vs others, blue color asterisk for Dh+ Cu 10_5min Vs others in wild type hCTR1 and above mentioned mutant hCTR1 variants. (G). Comparative analysis of normalized area fraction for mNeonGreen-hCTR1 puncta in same ROI of the cell against different copper treatments mentioned earlier [blue line represents wild type FLAG-hCTR1; green line represents ΔM1M2-FLAG-hCTR1; and dark brown line represents Δ30-FLAG-hCTR1].

### Membrane cholesterol is key in the regulation of CTR1 endocytosis

We established the critical role of copper-induced cluster formation as a pre-endocytic process in regulating CTR1 endocytosis. We found that membrane physical properties contribute significantly during several phases of Cu-induced regulatory endocytosis of CTR1.

Cholesterol is an essential lipid bilayer component ^48–51^. Studies by Biswas et al. have demonstrated that cholesterol depletion by methyl-β-cyclodextrin (MβCD) significantly remodels the plasma membrane’s mechanics by affecting pm tension ^52^. Kumar and Chattopadhyay demonstrated that intracellular sorting of the serotonin_1A_ receptor is altered upon mild cholesterol depletion and with increasing and acute cholesterol depletion complete inhibition of endocytosis of receptor is noticed.

In our coarse-grained MD simulations, we analyzed the presence of cholesterol around 0.6 nm of the proteins in the protomer and 9-mer oligomer states. We observed a significant increase in the number of cholesterols around the 9-mer oligomer compared to the protomer (Fig. 6A). Further, the spatial density analysis reveals that prominent cholesterol density is observed in the lower leaflet around the 9-mer oligomer at the curvature region (Fig.6B). This analysis hints towards a probable link of cholesterol with CTR1 oligomerization and membrane curvature. Taking this further, experiments were performed to understand the function of cholesterol on the CTR1 clustering and endocytosis.

**Figure: 6.**
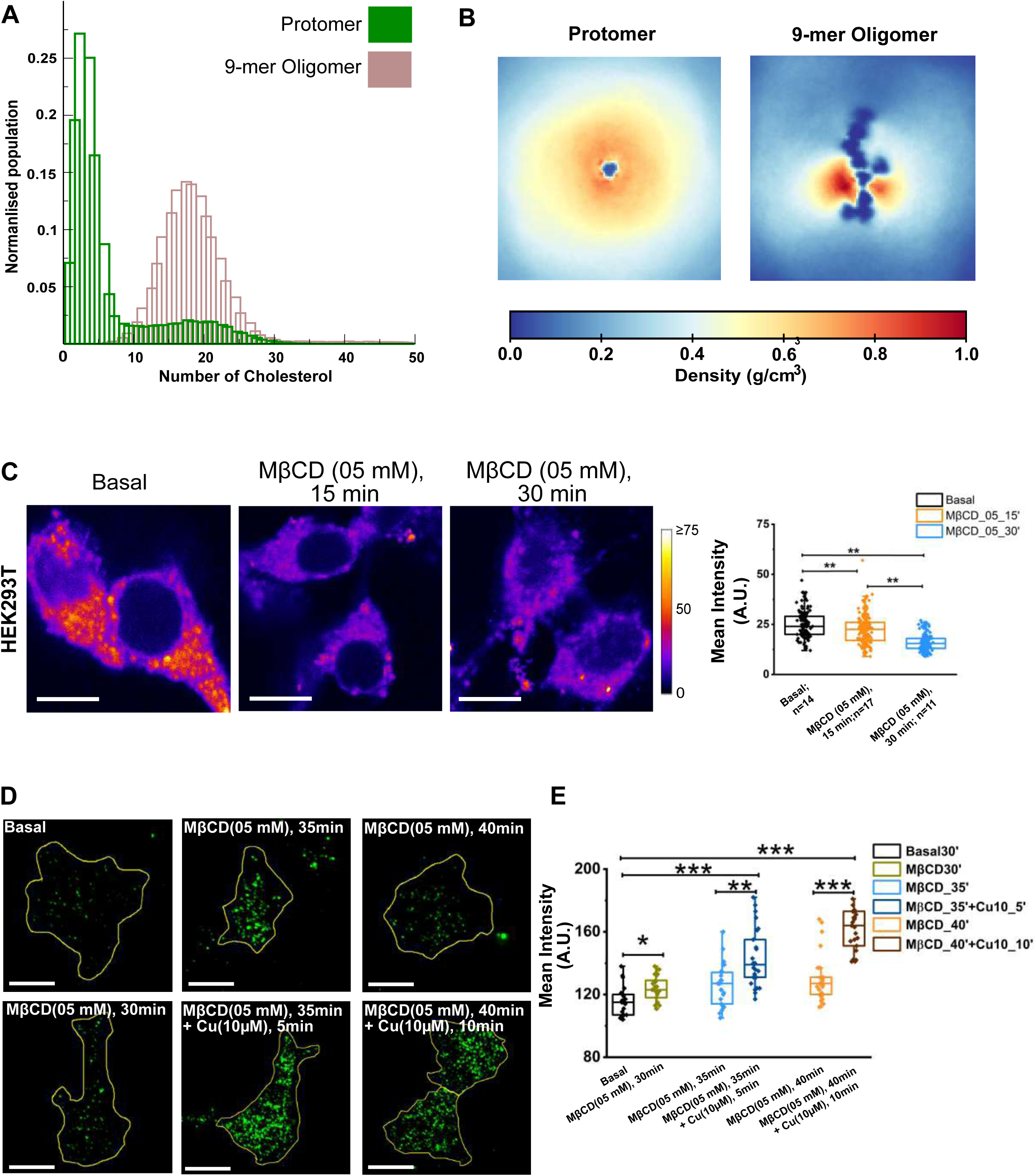
Membrane cholesterol is crucial for CTR1 clustering and endocytosis. A. Histogram from MD simulation studies depicts the cholesterols observed around the Protomer and 9-mer oligomer in the membrane. B. 2D map of the spatial density of cholesterol in the lower leaflet of the membrane around the protomer and the 9-mer oligomer analyzed through MD simulations. C. STED image of HEK293T cells stained with filipin-III for basal and MβCD (05mM; for 10min and 30min) treatment. Mean intensity plotted against filipin stain for the abovementioned treatments (Right panel). *N_ROI_* = 14, 17 & 11 respectively for untreated and the 10min and 30min MβCD treatments respectively. D. Representative TIRF image of FLAG-tagged hCTR1 (pseudo color green) expressed in HEK293T cells for basal, MβCD (05mM; 30min, 35min and 40 min) and MβCD along with copper treatment (10μM; 05min and 10 min). E. Comparative mean intensity profile of surface localization of hCTR1 for the abovementioned treatments (*N_ROIs_* = 20 for all experimental conditions). Statistical analysis was performed using Mann–Whitney U-test, ns denotes P value>0.05. * denotes P value <0.05** denotes P-value<0.001. Scale bars: 10μm.

We induced cholesterol depletion using MβCD (5mM) treatment for 30 minutes. Subsequently, the cells were stained with filipin, a fluorescent antibiotic that binds to cholesterol, and imaging was performed using confocal microscope. MβCD effectively disrupts filipin-bound cholesterol, as evidenced by a significant drop in mean intensity with time (Fig 6. C).

We further utilized TIRF microscopy to evaluate the surface retention of hCTR1 in the plasma membrane in cholesterol disrupted cell under excess extracellular copper conditions. We found that cholesterol disruption resulted in an increase of mean hCTR1 intensity at surface plasma membrane in a time dependent manner under excess extracellular copper level indicates delay in Cu-triggered CTR1 endocytosis (Fig 6. D & 6. E). Mentionable, though surface retention was higher, we did not observe any appreciable CTR1 cluster in cholesterol depleted cells treated with copper (Fig 6D). To summarize cholesterol depletion fails to cluster CTR1 in copper treatment and reduces the rate of Cu-induced CTR1 endocytosis, highlighting the crucial role of cholesterol in this regulatory process.

### CTR1 clusters recruit AP-2 and clathrin that primes it for endocytosis

We further investigated the downstream CME processes that is subsequent to copper-mediated CTR1 clustering. The Adaptor protein 2 (AP-2) is crucial for regulating Clathrin-Mediated Endocytosis (CME) by interacting directly with cargo through its mu (μ) subunit. Major AP-2 binding motifs found on the cargo molecules include the (a) di-leucine motif and the YXXL motif ^53,54^. Putative AP-2 binding domains were identified on the cytosolic loop of hCTR1, specifically between transmembrane domains 1 and 2, including motifs ^103^YNSM^106^ (YXXL) and di-Leucine motifs ^93^LL^94^ and ^133^LL^134^ (Fig.7A). During the pre-endocytic phase, induced by copper and Dh treatment, there was a noticeable increase in colocalization between hCTR1 and AP-2 as compared to basal conditions. (Fig.7B). We further utilized Stimulated emission depletion (STED) microscopy to gather a detailed view of CTR1-AP2 colocalization on the pm in copper treated and copper-depleted conditions. As compared to basal and copper depleted conditions, we recorded a significant increase in CTR1-AP2 colocalization in 10µM Copper treatment for 5 mins that further increase at 10 mins treatment (Fig 7C). We biochemically confirmed an increase in AP-2 and CTR1 interaction in copper-induces pre-endocytic state. AP-2 was slightly more abundant in the immunoprecipitated complex pulled-down with CTR1 antibody from copper treated cells (Fig 7D). The AP-2 adaptor complex links clathrin and cargo in clathrin-mediated endocytosis. Using immunofluorescence, we observed increased colocalization between hCTR1 and clathrin in pre-endocytic condition (Cu+Dh) and copper-induced endocytic conditions (only Cu) (Fig 7E). Notably, co-localization was appreciably higher when hCTR1 was arrested in the pre-endocytic stage. Immunoblot analysis showed that copper did not affect hCTR1 and Clathrin heavy chain expression levels (Suppl Fig 5). Immunoprecipitation with anti-hCTR1 antibody revealed the presence of clathrin in the pull-down complex when CTR1 was arrested at the pre-endocytic state, suggesting that copper treatment promotes clathrin recruitment in plasma membrane pits where hCTR1 aggregates (Fig 7F).

**Figure - 7.**
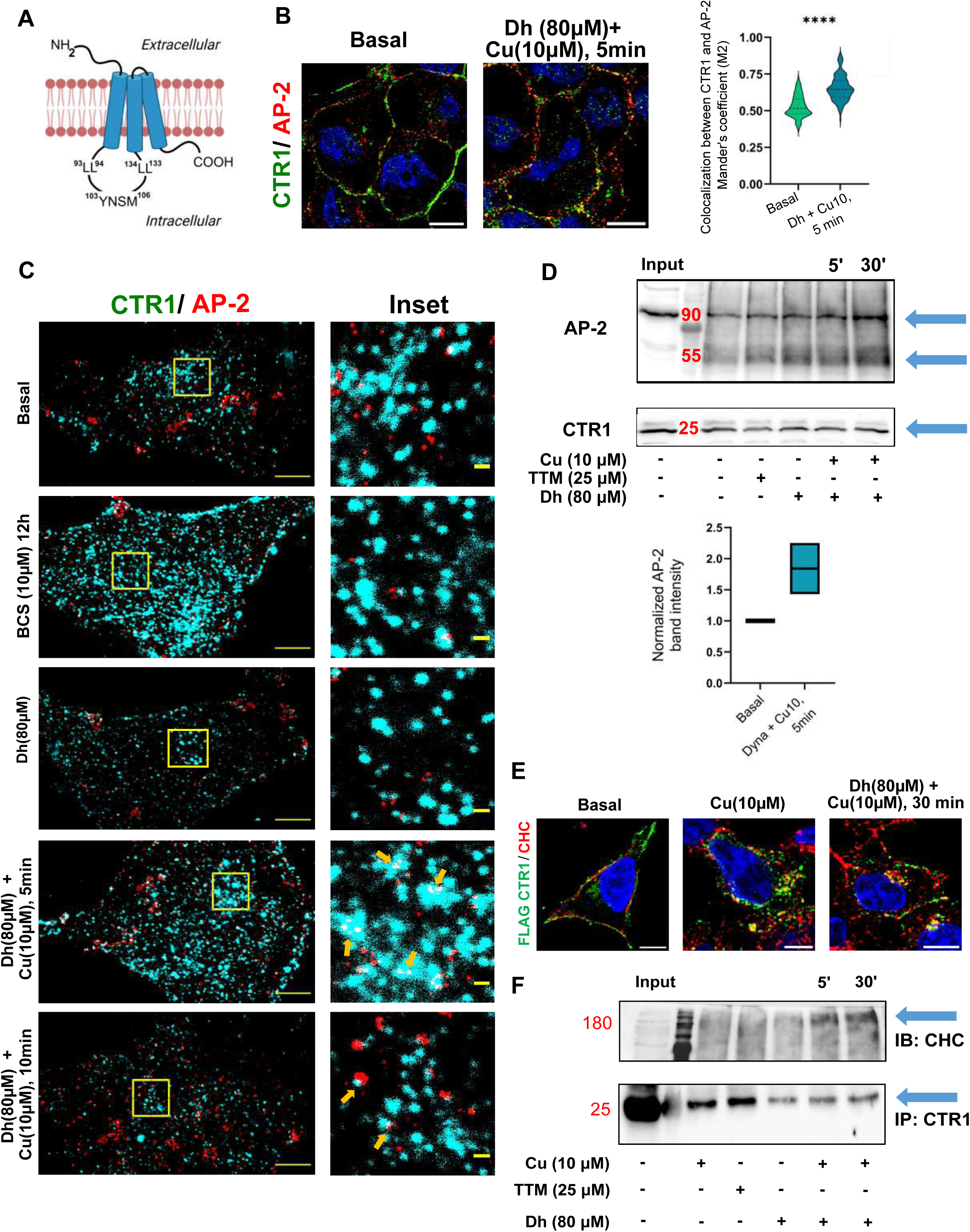
Copper treatment recruits the CME machinery to plasma membrane localized CTR1. A. Schematic illustration of the monomeric hCTR1 and putative AP-2 binding motifs in the cytosolic loop (Created in BioRender. Gupta, A. (2025) https://BioRender.com/t45r937) B. Localization of hCTR1 and AP-2 in HEK293T cells in basal (*left panel*) and dynasore (Dh) + Cu (10μM; for 5 min) treated condition (*right panel*); hCTR1 (*green*), AP-2 (*red*), DAPI (*blue*). The graph represents the comparison of Mander’s coefficient between hCTR1 and AP-2 at the above-mentioned treatment conditions; Violin plots represent the Mander’s colocalization coefficient (M2) for CTR1 colocalized with AP-2 under basal conditions (green) and following treatment with Dh + Cu10 for 5 minutes (blue). A significant increase in colocalization was observed upon treatment, indicating enhanced interaction or proximity of CTR1 and AP-2 (p < 0.0001, ****). Data were analyzed using Mann-Whitney U test, with two-tailed P value. C. STED images of the surface of HEK23T showing hCTR1 (green) and AP2 (red) localization in basal, Dh treated, bathocuproinedisulfonic acid disodium salt (BCS; 10µM for 12h), Copper (10µM; for 5 min and 10min with 20 min preincubation with 80µM Dh). Scale bars: 10µM D. Immunoblot with the copper and copper chelated (TTM) samples from co-immunoprecipitation (CO-IP) experiment; CO-IP was done with hCTR1 antibody and eluted samples were probed with Anti-AP-2 and Anti-hCTR1. In the graph, two independent experiments were analyzed. Bar graphs depicting normalized band intensity of AP-2. Band intensity of AP-2 were normalized against corresponding CTR1 band intensity and the values for the basal conditions were adjusted to one (1) for comparison. E. Immunofluorescent confocal microscopy images representing colocalization of hCTR1 (*green*) and Clathrin (*red*) (DAPI, blue) in basal, copper (10µM) treated and Cu (10µM, 30 min) in presence of Dh. F. Immunoblot from the coimmunoprecipitation experiment of hCTR1 and Clathrin Heavy Chain in photo-crosslinked cell samples using anti-CTR1 antibody. Cell treatment conditions included basal, copper, copper chelated, only Dh and copper treated for 5min and 30min in presence of Dh.

Fig 8 depicts the model that we propose based on our findings. To summarize, at the outset in basal copper, CTR1 is randomly distributed on the pm in a punctate fashion. With slight increase in copper, at early time points, CTR1 starts to form clusters facilitated by the Met-rich amino terminus of the protein. Membrane tension falls and fluctuation elevates possibly to accommodate these lateral movements and formation of the CTR1 clusters within the membrane. With further increase in copper and at later time points, once the cluster matures and CCPs starts forming, recruitment of AP-2 and Clathrin sets in; the endocytosis tug causes increase in membrane tension coupled with fall in membrane fluctuation. We coined this short window of events ‘*pre-endocytosis*’ that primes the CTR1 clusters for endocytosis, the process limits copper uptake and protects the cell from copper-induced toxicity.

**Figure - 8.**
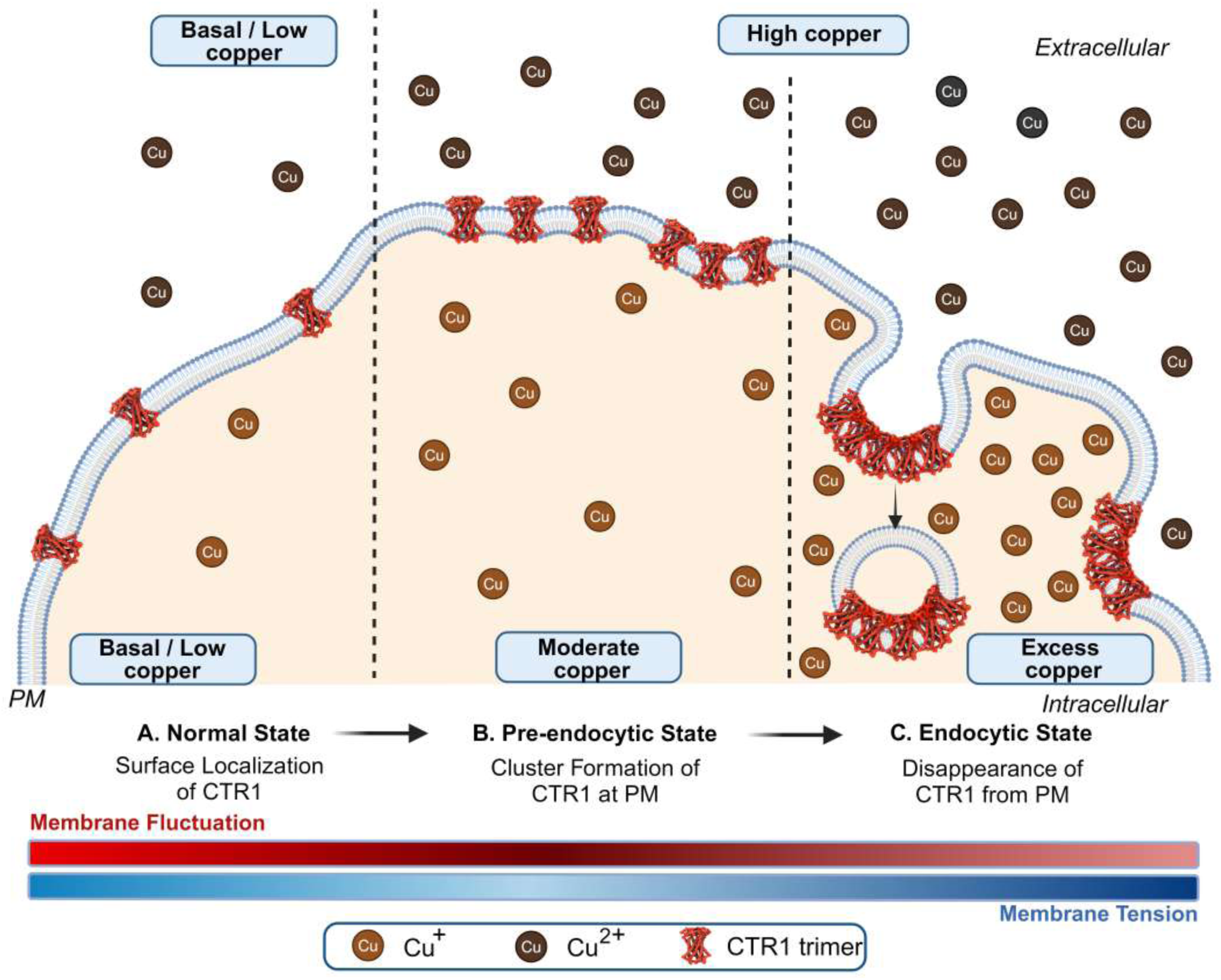
Model depicting Cu-induced CTR1 clustering and subsequent membrane property modulations. In basal/limited Cu condition, plasma membrane tension remains at a basal state and hCTR1s are sparsely distributed on PM, while continuing to import Cu into the cell. With increasing copper, sparsely distributed hCTR1 units start to appose on the PM and clusters are observed leading to the ‘pre-endocytic state’ of CTR1. This phase is marked by increase in membrane fluctuation and lowering of membrane tension. Next, as copper increases further, hCTR1 clusters localize on the clathrin-coated pits and membrane tension increases due to initiation of the membrane ‘tug’. This phase is marked by decrease in membrane fluctuation (Created in BioRender. Gupta, A. (2024) BioRender.com/i36d911).

## Discussion

Intracellular copper levels need to be tightly regulated as excess copper leads to cellular oxidative stress causing severe cellular damage. Tuning of cellular copper levels are primarily achieved at the levels of (a) copper import, regulated by CTR1 and (b) copper export controlled by the Cu-ATPases, ATP7A and ATP7B. We presented a mechanistic view of regulation of copper uptake by the mammalian copper transporter CTR1. The uniqueness of the finding lies in the fact that copper uptake by CTR1 is regulated by endocytosis that in turn is controlled by its copper-induced cluster formation on the pm. These clusters modulate membrane properties that primes CTR1 for endocytosis.

Channels and transporters rarely form clusters on the membrane to limit its ligand uptake. Generally, ion channels exhibit cluster formation to induce signal amplification and accelerate downstream response.

Copper selectivity by CTR1 lies in its TM domains. Studies by Bel-Tal’s group based on Cα-trace model of hCTR1’s TM domain supported by mutagenesis data revealed global movements of the channel associated with gating, selectivity, pH regulation, and overall conformational changes occurring during copper transport ^55^. The model uncovered that the TM-harboured Met^150^ and Met^154^ triads, by controlling gating and selectivity, regulate the passing of a single ion in every import cycle. The transport of the ion is facilitated by the directed rotational motion of the trimer. However, the model did not provide an understanding on how the protein ceases to uptake copper when the levels of the ion in cell reaches an optimum or saturation level.

We have previously reported that the copper-sequestering His-Met rich amino terminal of CTR1 regulate copper uptake. Deletion of the Met clusters or the proximal 30 residues harboring the 1^st^ His-Met residues not only abrogates copper-induced endocytosis of the protein but copper uptake competence is also compromised as determined by yeast-complementation assays ^12^. We could delineate a direct functional role of the amino terminal in copper uptake and subsequent CTR1 endocytosis. Our findings however, does not preclude the possible role of the cytosolic carboxy terminal that harbours the ^188^HCH^190^ motif in copper uptake.

Copper treatment induce CTR1 endocytosis. To gather a better understanding on the mechanism of copper uptake and CTR1 regulation, we asked how copper levels induce CTR1 endocytosis? While studying copper-induced endocytosis of CTR1, under confocal microscope, in live HEK293 cells, we serendipitously observed CTR1 clusters on the pm within a few minutes of adding 10µM or 50µM copper in the media. The clusters were short-lived and immediately vanished from the surface and underwent endocytosis. The observation attracted our attention as this copper-induced phenotype was specific for CTR1 and the pm marker Na,K-ATPase that colocalized with CTR1 under basal conditions did not exhibit such clustering. Up on delving into further detail to understand the mechanism of clustering using coarse-grained MD simulations and coupled IRM-TIRF microscopy, we found that CTR1 has an intrinsic property to form small puncta that enhances into larger clusters in presence of high extracellular copper (copper in the media). Presence of the extracellular amino terminal domain harboring the His-Met motif is indispensable to form copper-induced cluster and endocytosis. We however, due to experimental limitations could not quantify the minimum or maximum number of CTR1 channels that participate to form a endocytosis-inducing cluster. We calculated that cluster sizes varied from 100-250 pixels. Up on deriving the area of cluster in nm^2^, we found that cluster size varied from 422.25 nm^2^ to 1056.25 nm^2^ and large clusters were predominant and appreciably visible in presence of copper when endocytosis was stalled by dynasore.

Mechanistic understanding that implicates protein clustering at the pm with its endocytic regulation is inadequate. Paul et al ^56^ demonstrated that clustering of protein ectodomains triggers their endocytosis via a macroendocytic route, though that is independent of clathrin and dynamin. We hypothesized that cluster being a physical entity with an appreciable size and mass exerts localized mechanical stress on the plasma membrane. Indeed, we noticed that though copper, in general, does not affect plasma membrane properties, CTR1 clusters significantly alters local biophysical properties of the pm that facilitates endocytosis of the protein. We focused on determining the effect of CTR1 cluster on pm fluctuation and tension, two inversely related membrane mechanical properties that modulates clathrin mediated endocytosis.

Consistent with published data on mechanism of endocytosis, at early time points of copper treatment, we recorded a rise in membrane fluctuation that is indicative membrane addition or accumulation at sites on the pm that are being primed for endocytosis. Subsequently, membrane fluctuation falls accompanied with a rise in tension that demarcates the tug during the outset of formation of the CCPs. We named the window of ‘CTR1 clustering’ as its ‘pre-endocytic state’ as it preceded endocytosis and is a pre-requisite for endocytosis of the protein. Membrane tension settles back to basal levels up on completion of endocytosis.

Cholesterol is a key factor in tuning this pre-endocytic state. Interestingly, we noticed that loss of cellular cholesterol significantly slows down copper-mediated endocytosis of CTR1. At the same time, we failed to record formation of any CTR1 clusters on the membrane in presence of copper and dynasore. This data not only implicates cholesterol as a key ingredient in CME of CTR1 but also reinstates the fact that CTR1 clustering is a prerequisite of endocytosis.

Membrane clustering leading to endocytosis is typically observed in membrane receptors and is rarely observed in ion channels. The answer to why ‘reminiscent of a membrane receptor, ligand induced clustering occurs in CTR1 that leads to its subsequent endocytosis, probably lies in the fact that CTR1 used to act as a pm receptor for a now-extant paleoviruses CERV-1 and CERV-2. Studies from Beniasz’s group^57^ identified CTR1 as a receptor that was presumably used by CERV2 during its ancient exogenous replication in primates. Expression of human CTR1 was sufficient to confer CERV2 entry on otherwise resistant hamster cells. Interestingly, induction of CTR1 internalization of copper treatment inhibited CERV2 infection of human cells.

Our study provides a hitherto unknown mechanistic understanding of copper-induced regulation of the mammalian copper transporter CTR1. We show that copper induces clustering of CTR1 at the pm that precedes its endocytosis. Clustering significantly alters local mechanical properties of the membrane that facilitates CTR1 endocytosis that in-turn ensures optimal copper uptake in the cell and protect it from copper-induced toxicity.

## Methods

### Cell culture

HEK293T cells (ATCC) were cultured in Dulbecco’s modified Eagle’s medium (DMEM, #CC3004.05L, Cellclone) supplemented with 10% fetal bovine serum, 1× penicillin-streptomycin, and 1× amphotericin B. Cells were seeded in a T-25 flask (Thermo Scientific, #156367) and maintained by changing the medium on day 2 and splitting at 75–80% confluence on day 3 for passaging. For plasmid transfection, JetPrime reagent (#114-07, PolyplusTransfection) was used following the manufacturer’s instructions. The transfection mix (plasmid + JetPrime solution + buffer) was prepared in a sterile hood, incubated for 40 minutes, and added dropwise to cells in DMEM (−/−) medium. After 6 hours, the medium was replaced with complete DMEM (+/+), and cells were incubated for 16 hours to allow protein expression. Cell line was confirmed to be contamination-free.

### Plasmids and antibodies

Human CTR1 (hCTR1) was cloned into the p3XFLAG-CMV10 vector (Sigma, #E7658, gift from Dr. Rupasri Ain, CSIR-IICB) using HindIII and EcoRI restriction sites to generate an amino-terminal 3X FLAG-tagged construct. Amino-terminal mutations were introduced using the Q5® Site-Directed Mutagenesis Kit (NEB, #E0554) as per protocol, with primers designed accordingly and synthesized by GCC Biotech (West Bengal, India). For the generation of mNeon-tagged hCTR1, hCTR1, and mNeonGreen were PCR-amplified from lab plasmids, and the CMV backbone was amplified from the lyso20 mCherry plasmid (Addgene, #55073). PCR products were purified with the G-Bioscience Gel Purification Kit (G-Bioscience, #786-359-300P) and ligated using the NEBuilder HiFi DNA Assembly Kit (NEB, #E2621), following transformation into *E. coli* TOP10 cells, with successful clones confirmed by sequencing. Antibodies used included rabbit anti-FLAG M2 (CST, #14793), mouse anti-FLAG (CST, #8146), rabbit anti-CTR1 (Abcam, #ab129067), mouse anti-alpha Adaptin (Invitrogen, #MA3-061). Plasmid isolations were conducted using the QIAGEN Plasmid Mini Kit (QIAGEN, #27104).

### Dosage administration

For copper treatment, A stock solution (10 mM) of copper chloride (SRL) dissolved in water was used. HEK 293T cells were treated with 10μM and 50μM final concentration of copper chloride after 10 min, 30 min & 1hour at 37^0^C. For simulating the copper-chelated condition, a stock solution (2.5 mM) of TTM (Sigma #323446) dissolved in DMSO (Sigma #D2650) or 10 μM of bathocuproine disulfonate (BCS; Sigma) dissolved in distilled water has been used^58^. To inhibit all dynamin-dependent endocytic pathways, we incubated cells with Dynasore hydrate (Dh) 80 μM; Sigma) in serum-free media for 20 min ^59–61^ followed by 10μM copper treatment for different time courses. 5 mM Methyl-ß-cyclodextrin (Sigma-Aldrich) was used in serum-free media for 50 min to deplete Cholesterol ^62^.

### Immunofluorescence

For HEK-293T cells, glass coverslips were affixed to the bottom of a 24-well plate (SPL, #30024) for cell cultivation. Upon reaching 60-70% confluency, HEK-293T cells were subjected to transfection and /or subsequent treatments. Following the designated treatment duration, the cells were rinsed 2-3 times with ice-cold PBS buffer, with each wash lasting 2 minutes, to eliminate remaining growth media and dead cells. The cells were subsequently fixed using an ice-cold 1:1 solution of acetone and methanol for 20 minutes at (−20°C). The cells were rinsed 3 to 4 times to eliminate the fixative solution. Subsequently, blocking and permeabilization were conducted using a blocking solution of 3% BSA in PBSS (0.075% saponin in PBS) for 3 hours at room temperature. Primary antibodies were administered to the cells in a 1% BSA in PBSS solution and incubated for 2 hours at room temperature. After that cells were washed three times (each wash lasting 5 minutes) with PBSS, and appropriate secondary antibodies were applied and incubated for 2 hours at room temperature. Following the incubation, the cells were gently washed with PBSS (three times, each wash lasting 5 minutes) and PBS (twice, each wash lasting 5 minutes), and finally, the cells were mounted using Fluoroshield with DAPI mounting medium.

### Confocal microscopy

All images were obtained using a Leica SP8 confocal platform with an oil immersion 63X objective (NA 1.4) and deconvoluted with Leica Lightning software. For a series of experiments, we maintained consistent image acquisition settings, including pinhole, zoom, laser power, emission gain, and scan speed.

### Image analysis

Image analysis was conducted in batches using ImageJ software. For colocalization analysis, the *Colocalization_Finder* plugin was utilized. Regions of interest (ROIs) were manually delineated on the optimal z-stack for each cell. Manders’ colocalization coefficient (MCC) was calculated to quantify colocalization. For cluster analysis, the *Particle_Analyzer* plugin was employed. Images were converted to binary format, and thresholding was applied using the default settings. Macro codes used in ImageJ analysis for chapters one and two are available at https://github.com/saps018/hCTR1-N-term/tree/main/colocalization and https://github.com/saps018/CTR1-clustering respectively. Data visualization and statistical analysis were performed using GraphPad Prism (version 10.3.1). Graphs, including bar and scatter plots, were generated to represent data, with statistical significance determined by t-tests and ANOVA where appropriate.

### Immunoblotting

HEK-293T cells were cultured in 60 mm dishes (Genetix, #GX-02060) to 70–80% confluency. Following specific treatments and transfection, cells were placed on ice, and media were discarded. Cells were washed twice with ice-cold PBS, then scraped into ice-cold PBS using a cell scraper (Tarsons, #960052) and collected by centrifugation at 3,500 RPM for 5 minutes. Cell pellets were resuspended in RIPA lysis buffer with gentle pipetting, then incubated on ice for 60 minutes with intermittent mixing. The lysates were sonicated (5 pulses, 5 s/pulse, 100 mA) with 30-second intervals on ice. After centrifugation at 3,000g for 10 minutes at 4°C, the supernatant was collected for protein quantification via the Bradford assay (Sigma-Aldrich, #B6916-500ML).

Proteins were mixed with Laemmli buffer and separated by SDS-PAGE (10% gel). Proteins were transferred onto nitrocellulose membranes (BioRad, #1620112) using a semi-dry transfer system (BioRad, Trans-Blot SD) at 15V. Transfer times were adjusted based on the target protein size. After blocking with 3% BSA in TBST (1x TBS with 0.01% Tween-20) for 3 hours at RT, primary antibody incubation was performed overnight at 4°C. Membranes were washed thrice with TBST (10 minutes each), incubated with HRP-conjugated secondary antibody for 2 hours at RT, and washed again with TBST and TBS. Signal detection was performed using ECL reagent (BioRad, #170-5060) and visualized on a ChemiDoc system (BioRad).

### Co-immunoprecipitation

Cells were cultured in 60-mm dishes. Following treatment, they were placed on ice and washed twice with ice-cold PBS. Cells were scraped in 1 mL of PBS, centrifuged at 3500 rpm for 3 minutes, and the supernatant was discarded to collect the dry cell pellet. Pellets were resuspended in 100-120 μL CHAPS lysis buffer (50 mM Tris-Cl pH 7.5, 150 mM NaCl, 0.1 mM CHAPS, 1x protease inhibitor cocktail, 1x PhosSTOP, 1 mM sodium orthovanadate, and 1 mM EDTA) and incubated on ice for 30 minutes with gentle pipetting. Lysates were then homogenized by passing through a 26G syringe (50 strokes) and incubated on ice for an additional 15 minutes. The lysates were centrifuged at 8000 rpm for 15 minutes at 4°C, and the supernatant was collected. For bead preparation, 75 μL of Protein G magnetic beads were washed three times with binding/wash buffer (20 mM Na2HPO4, 150 mM NaCl) using a magnetic rack, and resuspended in 75 μL of the same buffer. To each tube, 2 μL of CTR1 antibody (abcam, #ab129067) was added, and the beads were incubated with gentle rotation at room temperature for 1 hour. The beads were washed three times with wash buffer, then incubated twice with 0.2 M triethanolamine, pH 8.2. Crosslinking was initiated by adding 1 mL of freshly prepared 20 mM dimethyl pimelimidate (DMP) solution in 0.2 M triethanolamine, followed by incubation at room temperature for 30 minutes with gentle rotation. The reaction was quenched by resuspending the beads in 50 mM Tris-Cl, pH 7.5, and incubating for 15 minutes. After washing three times with PBS, pH 7.4, antibody-cross linked beads were incubated with cell lysates overnight at 4°C with rotation.

Following incubation, the supernatant was collected to assess unbound fractions, and beads containing the bound proteins were washed three times with PBS. Elution was performed using 100 μL of 1x LDS sample buffer containing 10% β-mercaptoethanol, heated at 95°C for 10 minutes. Eluted samples were collected using the magnetic rack and prepared for immunoblotting.

### Total Internal Fluorescence Microscopy (TIRFM) Imaging

Transfected HEK293T cells were exposed to MβCD alone or in combination with copper for various durations (0-60 minutes) and fixed with 4% PFA. Fixed cells were subjected to imaging using TIRFM. TIRFM was performed on an inverted microscope (IX81, Olympus Corporation, Japan) using 100× TIRF objective (N.A 1.49) and a CMOS camera (ORCA-Flash 4.0, Hamamatsu Photonics, Japan) with a 1 pixel=65 nm. **Suppl. Fig-1. A** illustrates the light path in TIRF. TIRF images were captured using PSS lasers of 488 nm wavelengths and at penetration depth of 70 nm. For all live cell imaging cells were maintained at 37°C throughout by using an onstage as well as cage incubator (Okolab, Italy).

### Interference Reflection Microscopy

An inverted microscope (Nikon, Ti-E eclipse, Tokyo, Japan) with adjustable field and aperture diaphragms, 60x Plan Apo (NA 1.22, water immersion) with 1.5x external magnification, 100 W mercury arc lamp, (546 ± 12 nm) interference filter, 50:50 beam splitter and CMOS (ORCA Flash 4.0 Hamamatsu, Japan) camera were used for IRM. Fast time-lapse images of cells were taken at 20 frames per second and total of 2048 frames were captured. Suppl. Fig. 4A, illustrates the interference reflection in IRM. Calibration, identification of FBRs and quantification of fluctuation amplitude (SD_time_) and tension were done previously reported^45^. Briefly, the membrane height fluctuation was calculated by converting the intensity profile into the height profile obtained from the IRM images. For intensity change (ΔI) to height change (Δh) conversion, a 60 μm polystyrene bead was imaged at different exposure times (Suppl. Fig. 4B). We also chose regions that lie between the maximum intensity and minimum intensity of the first branch of intensity profile of the bead so that any repetition of the same intensity can be avoided. The intensity plot profile that obtained from bead images has periodicity (intensity profile of bead) and shows repetition. To avoid misinterpretation due to the degeneracy, for analysis, only pixels falling on the first branch (or ∼ 100 nm of coverslip) were selected as regions of interest (ROIs) and termed as First Branch Regions (FBRs) (Suppl. Fig. 4B). The standard deviation (SD) of the relative height from a pixel-wise time series of membrane height fluctuations was termed SD_time_. Averaging pixel-wise measurements within FBRss, provided the FBR-wise data (Figure 2. E, Yellow small boxes within zoom in IRM image of HEK 293T cell). From all over the cell, the membrane that lies within 100 nm height from the coverslips (excluding the nuclear region) is utilized to get our FBR-wise analysis, which provides local information. Measurements on FBRs were averaged over single cells to provide cell–wise statistics. Further, we compared these parameters between basal and copper-treated cells to find the impact of copper-induced hCTR1 endocytosis on membrane mechanics. HEK293T cells were treated with copper (10 or 50µM) for various time (0-60 minutes) periods and subjected to IRM and for all live cell imaging cells were maintained at 37°C throughout by using an onstage incubator.

### TIRF-IRM Microscopy

For observing the CTR1 localization upon copper treatment along with membrane tension, correlative TIRF IRM imaging was performed on an inverted microscope (Nikon, Ti-E eclipse, Tokyo, Japan) using 100X objective (NA 1.40) with the above-mentioned IRM specifications, whereas TIRF images were captured using coherent OBIS lasers of 488 nm. For this, transfected HEK 293T cells were treated with 10μM conc. of copper chloride for 10 min followed by 10 min post incubation with TTM. Imaging involved acquisition of sequential TIRF (300 frames, 200 ms exposure), IRM (2048 frames, 20 fps) and TIRF (300 frames, 200 ms exposure) images following single fields.. To study the CTR1 localization and membrane tension at the pre-endocytic phase, transfected HEK 293T cells were preincubated with dynasore hydrate (Dh) and treated with 10μM conc. of copper chloride for 10 min, and simultaneous images were acquired using the IRM-TIRF setup. The first frame of the TIRF movie obtained after the IRM movie was used for the correlative analysis. Comparison of the TIRF movies before and after the IRM movie was used to verify the stability of the observed fluorescent pattern over the 1-2 min imaging period.

### STED imaging

HEK293T cells were treated with 10μM Copper treatment preincubated with or alone Dh and compared to basal and copper depleted (10 µM; BCS treatment for 2hr) conditions. After treatment washing with ice-cold PBS (2 × 2 min), cells were fixed with Acetomethanol (1:1, v/v; while using hCTR1 antibody) for 20min at room temperature (RT), followed by 20 min incubation with 50 mM ammonium chloride (SRL) in PBS. Next, the cells were washed with PBS, and blocking was performed in 3% Bovine Serum Albumin (BSA, SRL #85171) in PBSS (0.075% saponin in PBS) for overnight at 4^0^C. Primary antibody treatment was performed (hCTR1, Clathrin heavy chain and AP2-µ subunit) at 1: 200 dilutions in 3% BSA in PBSS for 3 h at RT followed by PBSS washes (3 × 5 min). After that, incubation with the respective secondary antibodies (Alexa 568; (Abcam) and STAR Red; (Abberior) conjugated secondary antibodies were used for imaging CTR1 and AP2, respectively) was used at 1:1000 dilution for 2 h followed by PBS washes (3 × 5 min) and mounted with the MOWIOL (DABCO, Sigma, D2522). A Facility Line system with an Olympus IX83 microscope (Abberior Instruments) was used for STED imaging. Abberior auto alignment sample was utilized for alignment of the STED and confocal channels. 15 nm pixel size was maintained during imaging. A pulsed STED line at 775 nm was used for depletion.

### Metabolic labelling with photo-amino acids

Photo-reactive amino acid crosslinking was conducted in accordance with the manufacturer’s guidelines (Pierce Biotechnology). Initially, HEK-293T cells were cultivated in standard complete growth media (DMEM) as previously outlined on 60 mm tissue culture dishes. Upon reaching 60-70% confluency, the standard complete growth media was removed, and leucine- and methionine-free Dulbecco’s modified Eagle’s limiting medium (DMEM-LM, Ref. #30030, Lot #UG2807111, Thermo Scientific) was supplemented with 10% dialyzed serum (Dialyzed FBS one shot, TM, SKU no. #A3382001, Thermo Fisher), 1% penicillin-streptomycin, L-Photo-leucine (Ref. #22610, Lot No. #UC273699), and L-photo-methionine (Prod. #22615, Lot No. #UD273716) to achieve final concentrations of 4 mM and 2 mM, respectively. Cells were maintained in specialized media for 24 hours, during which particular therapies were administered. All treatments were conducted in Dulbecco’s modified Eagle’s limiting medium devoid of leucine and methionine (except the 10% dialyzed serum). Following a 24-hour incubation, the cells were removed from the incubator and placed on a cool surface, where they underwent two washes with ice-cold PBS to eliminate residual media and dead cells. Subsequently, 1 ml of cold PBS was gently poured to the dish, ensuring that the cell attachments were undisturbed and that the whole surface of the attached cells was covered. Subsequently, cells were irradiated with a UV lamp at 350 nm for 20 minutes at room temperature from a distance of 4 cm; during this interval, the incorporated photo-leucines and photo-methionines were cross-linked with adjacent molecules. Subsequently, the cells were fixed for immunostaining or resuspended in PBS solution and prepared for immunoblotting as previously outlined.

### Molecular Dynamics Simulations

Coarse-grain molecular dynamics simulations were performed with a functional trimer unit or protomer of hCTR1 proteins embedded in model membranes to understand the effect of proteins and protein clusters on the lipid bilayer.

#### System setup and simulation parameters

The initial atomistic protomer structure of hCTR1 was obtained based on the electron crystallography structure provided by Professor Vinzenz Unger, Department of Molecular Biosciences, Northwestern University ^8^, which has been previously to study the molecular mechanism of the copper binding at the N-terminal domain of hCTR1 ^12^.

#### Coarse grain system setup and parameters

The atomistic structure was energy minimized and mapped to the coarse grain using the martinize2 script for the Martini3 force field ^63,64^ with the Martini Gō model ^65^. The dissociation energy of the Lennard-Jones potential value 12 kJ/mol was used to represent Gō potentials. The functional unit trimer hCTR1, i.e., protomer, was embedded in the lipid bilayer with lipid composition using insane.py script ^66^. The lipid composition used for all simulations setup was upper Leaflet with 12% POPC, 16% POPE, 10% POPS, and 14% CHOL; Lower Leaflet with 28% POPC, 4% POPE, and 16% CHOL. Molecular dynamics simulations were performed using the Martini3 force field with the GROMACS simulation package (version 2021) ^67^. The system temperature was maintained at 300K by coupling to a V-rescale thermostat with a 1.0 ps coupling constant ^68^. A pressure of 1 bar was maintained using the semi-isotropic pressure coupling scheme with the Parrinello-Rahman barostat with coupling constant 1.0 ps and compressibility 3×10^−4^ bar^−1 69^. The non-bonded interactions were treated with a reaction field from 0.0 to 1.1nm for the coulomb interactions and 0.0 to 1.1 nm for the LJ interactions. A time step of 20 fs was used. Each system was solvated with water and neutralized with NA+ and CL-ions.

Simulations were performed for hCTR1 protomer with varying copy numbers in the membrane. A representative snapshot is shown figure 1.C and 1.D. We considered the following setups/systems with given copies hCTR1 protomer: 1) Only membrane: membrane + (0 protomer) hCTR1; 2) Protomer: Membrane + (1x protomer) hCTR1; 3) Multimer: Membrane + (9x protomer) hCTR1. Further, to understand the effect of these large clusters and the single oligomer (1 cluster of all nine hCTR1 protomers (9-mer)) on the membrane; 4) Oligomer: Membrane + (9-mer oligomer) hCTR1 in 1 cluster: This system was obtained by performing a biased simulation by pulling the 2 clusters of the hCTR1 (i.e. 4/5-mer clusters) observed in the multimer system simulations. Once the single oligomerized state was observed, again the equilibration of 2 microseconds was performed, and independent replicate simulations of the oligomer were performed. Standard parameters corresponding to the coarse-grain Martini simulations were used. For all the systems, simulations were performed in replicates of 5 for extended microseconds. The membrane, protomer (1x protomer), multimer (9x protomer), and oligomer (9-mer oligomer) systems were simulated for 225 μs, 182 μs, 455 μs, and 54 μs, respectively. The analysis was done on the last 10 microseconds of all simulation replicates.

### Analysis

All the simulations were analyzed using the inbuilt GROMACS tools and in-house scripts. Simulation snapshots and structures were rendered using VMD ^70^. For membrane thickness and curvature calculation, the GROMACS-compatible module g_lomepro was used ^71^.

#### Membrane tension

We simulated all the systems at NP_z_T ensemble to calculate membrane tension ^72^. Simulations were performed for 100 ns for all systems using the well-equilibrated conformations from each system type. The time-averaged membrane tension was calculated using the formula:

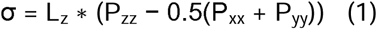

Here, L_z_ is the membrane thickness, P_xx_, and P_yy_ are the tangential components of the pressure tensor, and P_zz_ is the normal component of the simulation box.

#### Bending modulus

We used an in-house script to calculate bending modulus ^73^. The bending modulus (S(q)) of lipid membranes can be determined from a Fourier analysis of the height fluctuations (h) in the membrane of area (A). The following equation describes this relationship:

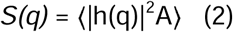

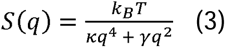

Where, κ is the bending rigidity, γ is the membrane tension, k_B_ is the Boltzmann constant, T is the temperature, and q is the wave number. The observed values were averaged over replicates and plotted as a bar graph with standard errors.

## Abbreviations

CTR1: Copper Transporter-1
MD: Molecular Dynamics
TIRF: Total Internal Reflection Fluorescence Microscopy
IRM: Interference Reflection Microscopy
CME: Clathrin Mediated Endocytosis

## Acknowledgement

This work was supported by DBT-Wellcome Trust India Alliance Fellowship (IA/I/16/1/502369) and Core Research Grant (CRG/2021/002150) from SERB, Department of Science and Technology (DST), Government of India STARS-2 Grant (2023-0210) from Ministry of Education, Govt. of India and IISER K intramural funding to A.G, DBT Builder project (BT/INF/22/SP45383/2022) to B.S and A.G; and Wellcome Trust/DBT India Alliance fellowship (grant number IA/I/13/1/500885)

S.K. and S.M were supported by pre-doctoral fellowship from Council of Scientific and Industrial Research and S.N. by post-doctoral fellowship from Department of Biotechnology, India. S.K.C was supported by post-doctoral fellowship from IISER Kolkata. T.G. was supported by CEFIPRA (grant number 6303-1) and IISER Kolkata. We thank IISER-Kolkata for providing us the support with the Central Microscopy Facility.

## Author contributions

Conceptualization: A.G., B.S. and D.S.; Methodology: A.G., B.S and D.S; Software: B.S, D.S and S.M.; Formal analysis: S.N., A.G., B.S and D.S.; Investigation: S.K.C, S.K., T.D., S.N. and T.G.; Resources: A.G., B.S and D.S.; Data curation and analysis: B.S., D.S, A.G., T.D., and S.M; Writing - original draft: A.G., B.S., D.S.,; Writing - review & editing: A.G., B.S., D.S., S.N., S.K.C. and T.D.; Project administration: A.G., B.S. and D.S.; Funding acquisition: A.G., B.S. and D.S.

---

**Supplementary Figure - 1.**
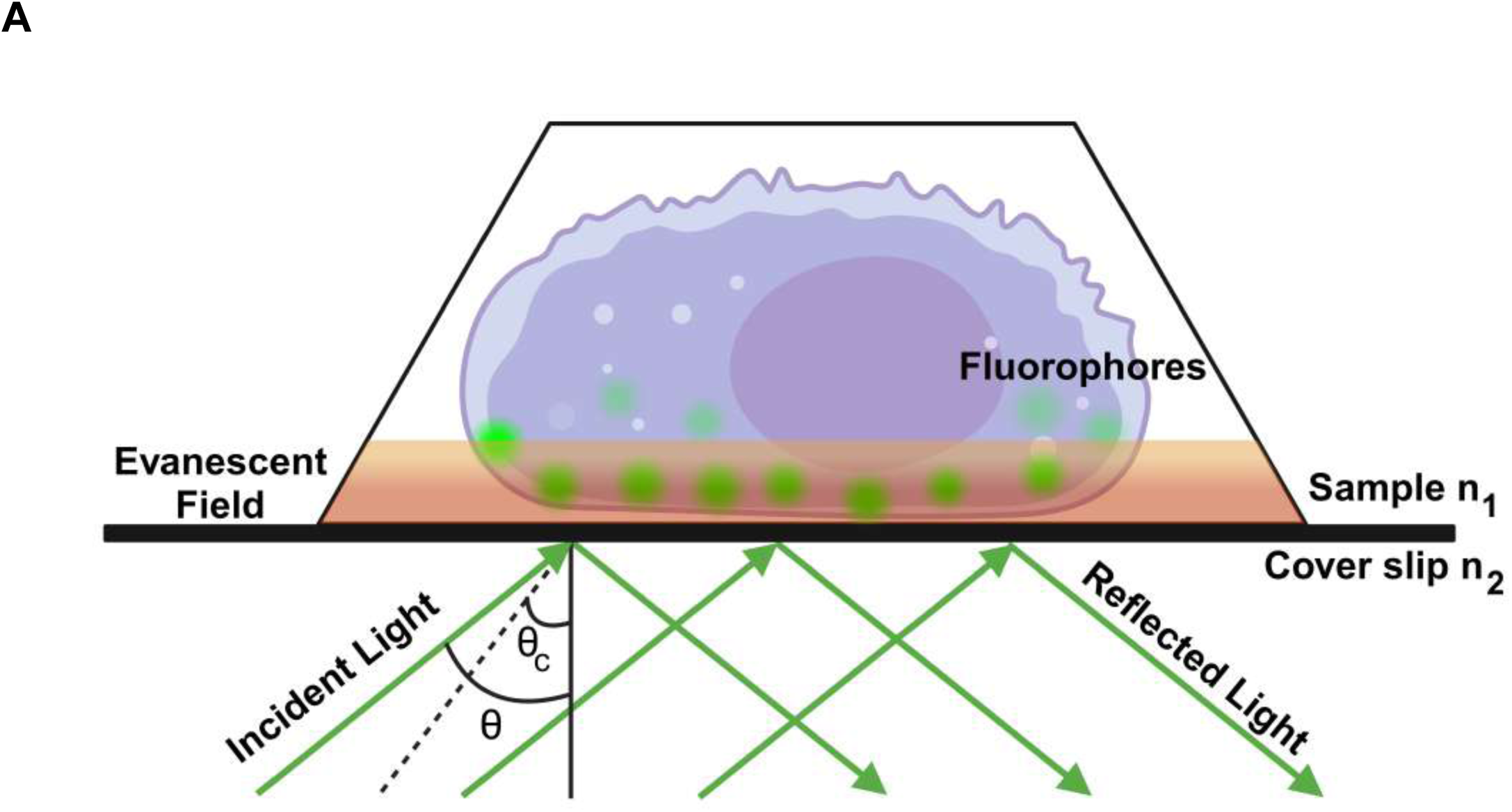
Principle of TIRF imaging. **A.** Representative diagram depicting the principle of Total Internal Reflection Fluorescence Microscope (TIRF).

**Supplementary Figure – 2.**
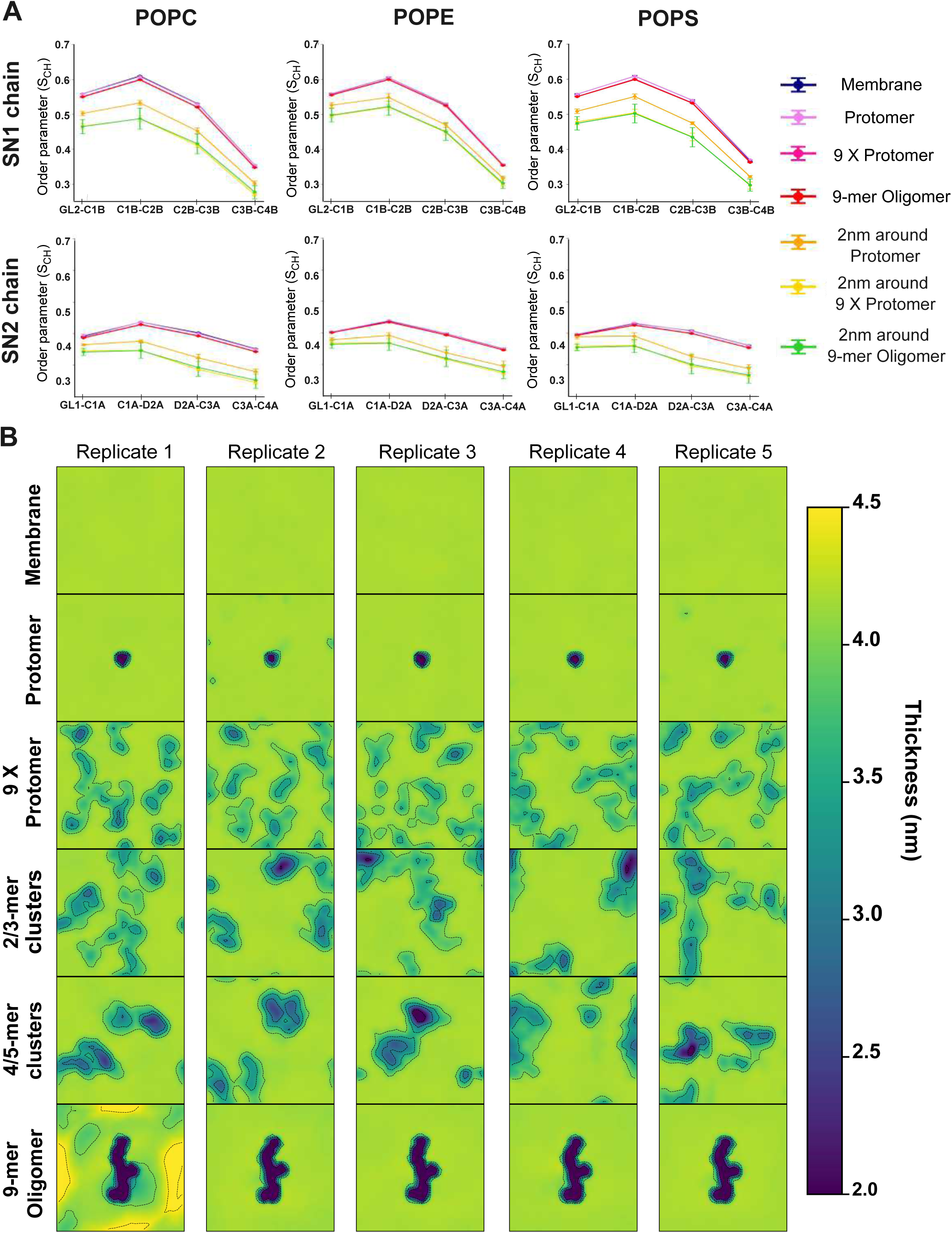
Modulation of membrane properties observed in Molecular Dynamics simulations. A. Lipid order parameters for acyl chains membrane lipids (POPC, POPE, and POPS are shown in the side panel) in the presence of different oligomeric states of hCTR1 and only the membrane. The parameter was calculated with standard error across replicates for all lipids and lipids at 2 nm around the hCTR1 proteins. The values reflect differences in acyl chain ordering between lipid types in the system. B. Membrane thickness across different oligomeric states of hCTR1, calculated for five replicates. The profiles indicate membrane perturbation effects within local regions surrounding different oligomers.

**Supplementary Figure – 3.**
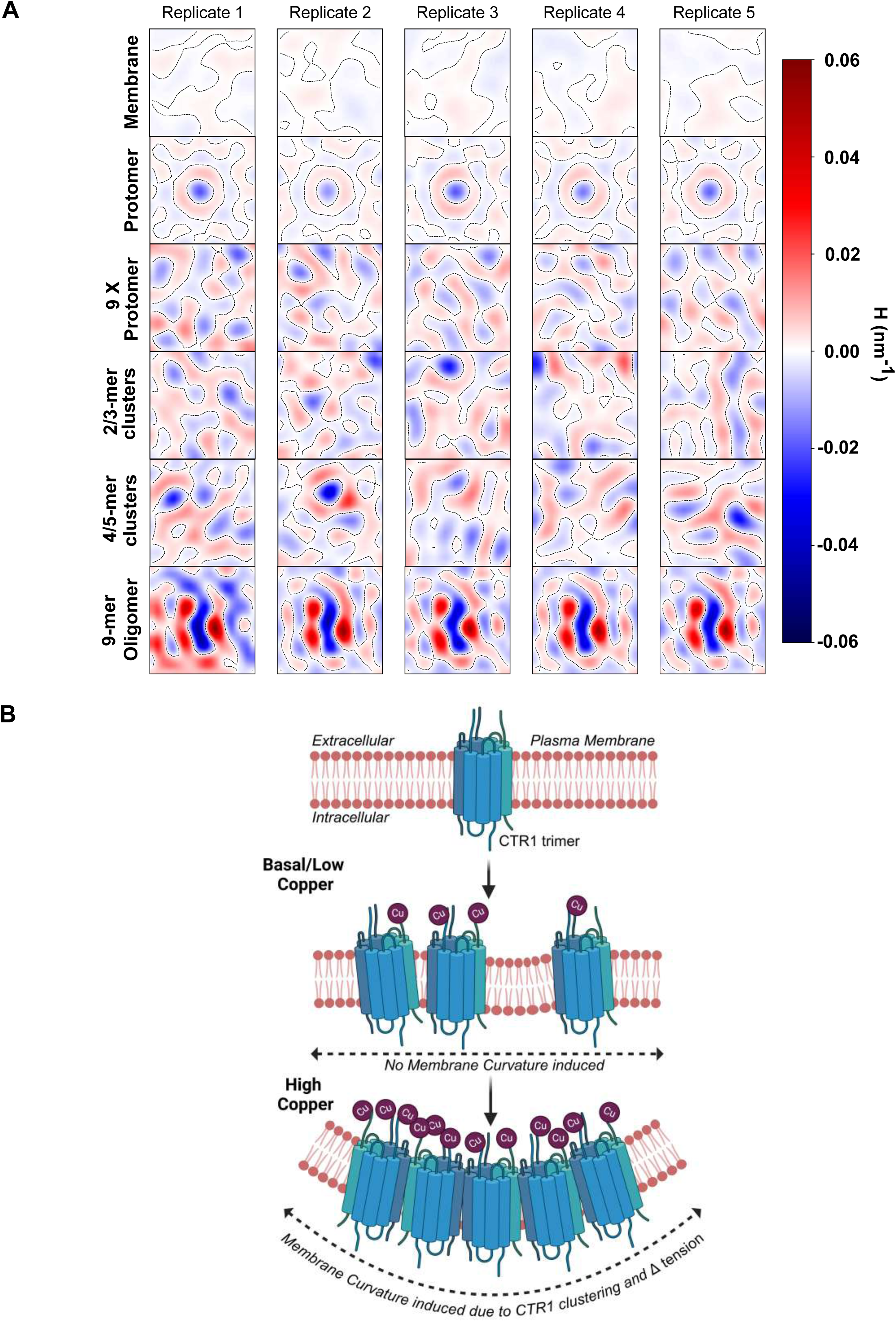
A. Membrane curvature was calculated for lower leaflets across different oligomeric states of hCTR1 and calculated for five replicates. The profiles indicate membrane curvature observed around different oligomeric states of the hCTR1. B. Hypothetical model deciphers CTR1 cluster-induced membrane curvature and subsequent alteration in Tension during CTR1 endocytosis regulation at elevated copper levels (Created in BioRender. Gupta, A. (2025) https://BioRender.com/n02y773)

**Suppl. Figure – 4.**
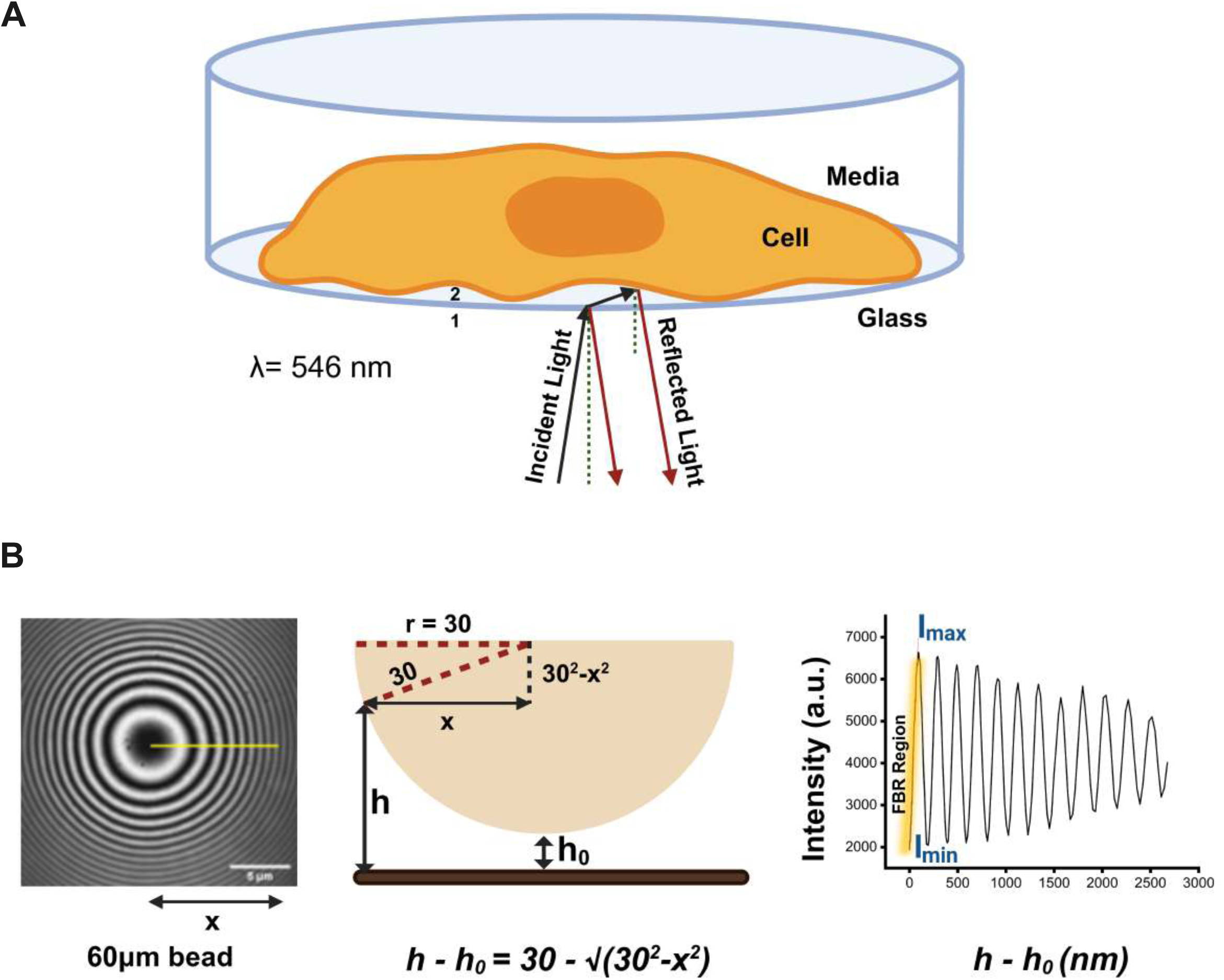
Principle of IRM imaging. A. Schematic diagram of Interference Reflection Microscope (IRM) set up (*Created in BioRender. Gupta, A. (2025)* https://BioRender.com/y57p602) B. Typical IRM image of 60μm polystyrene bead with a yellow radial line used for calibration to detect basal membrane height fluctuation within ∼ 100 nm from the coverslip (left). Schematic representation of a bead showing relative height from the coverslip, (Middle) A typical line profile plot of Intensity vs relative height. Red line shows linear fitted region which is taken as First Branch Region (FBR) for analysis and intensity minima (Imin) and intensity maxima (Imax) are denoted in blue color (Right) (*Created in BioRender. Gupta, A. (2025)* https://BioRender.com/e02p715).

**Suppl. Figure – 5.**
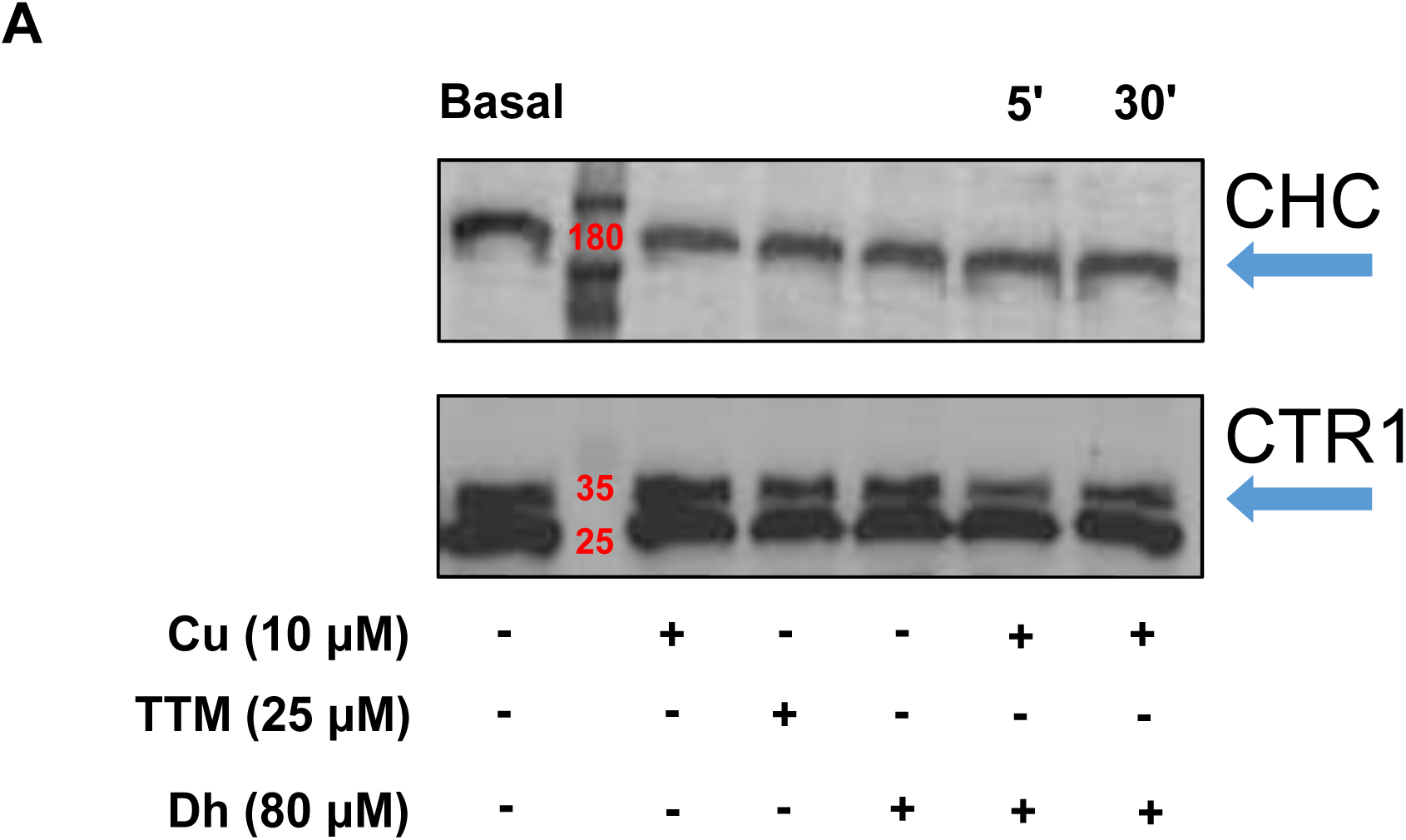
CTR1 and CHC abundance are not regulated by Copper. A. Immunoblot image showing the effects of copper (Cu), tetrathiomolybdate (TTM), and Dynasore (Dh) treatments on the levels of CHC and CTR1 proteins. Cells were treated with Cu (10 μM), TTM (25 μM), Dh (80 μM) and Cu, in the presence of Dh (80 μM) for the indicated time points (5 minutes and 30 minutes). CTR1 and CHC protein bands are marked by arrows, with molecular weights denoted in kilodaltons (kDa). “+” indicates the presence of the treatment, and “-” indicates its absence.

